# Pathogens, parasites, and parasitoids of ants: a synthesis of parasite biodiversity and epidemiological traits

**DOI:** 10.1101/384495

**Authors:** Lauren E. Quevillon, David P. Hughes

## Abstract

Ants are among the most ecologically successful organisms on Earth, with a global distribution and diverse nesting and foraging ecologies. Ants are also social organisms, living in crowded, dense colonies that can range up to millions of individuals. Understanding the ecological success of the ants requires understanding how they have mitigated one of the major costs of social living-infection by parasitic organisms. Additionally, the ecological diversity of ants suggests that they may themselves harbor a diverse, and largely unknown, assemblage of parasites. As a first step, we need to know the taxonomic and functional diversity of the parasitic organisms infecting ants. To that end, we provide a comprehensive review of the parasitic organisms infecting ants by collecting all extant records. We synthesize major patterns in parasite ecology by categorizing how parasites encounter their ant hosts, whether they require host death as a developmental necessity, and how they transmit to future hosts.

We report 1,415 records of parasitic organisms infecting ants, the majority of which come from order Diptera (34.8%), phylum Fungi (25.6%), and order Hymenoptera (25.1%). Most parasitic organisms infecting ants are parasitoids (89.6%), requiring the death of their host as developmental necessity and most initially encounter their hosts in the extranidal environment (68.6%). Importantly, though most parasitic organisms infecting ants only need a single host to complete their life cycle (89.2%), the vast majority need to leave the nest before transmission to the next ant host can occur (88.3%), precluding ant-to-ant transmission within the nest. With respect to the host, we only found records for 9 out of 17 extant ant sub-families, and for 82 out of the currently recognized 334 ant genera. Though there is likely bias in the records reported, both host and parasite ecological traits and evolutionary histories underlie the pattern of ant-parasite association reported here. This work provides a foundation for future work that will begin to untangle the ecological drivers of ant-parasite relationships and the evolutionary implications thereof.

## 2. Introduction

Ants (Hymenoptera: Formicidae) are among the most ecologically successful organisms on Earth (Hölldobler & Wilson, 1990; Wilson, 1990; Schultz, 2000). They account for a significant proportion of total insect biomass (Beck, 1971; Fittkau & Klinge, 1973; Tobin, 1995), and are important constituents of ecological communities (Schneirla, 1971; Hölldobler & Wilson, 1990; Folgarait, 1998). Ants inhabit almost every terrestrial ecosystem (Wilson & Taylor, 1967; Hölldobler & Wilson, 1990), have diverse nesting and foraging ecologies (Hölldobler & Wilson, 1990), and are speciose, with over 15,000 extant species in 334 genera (Bolton, 2018).

Modern ants (Hymenoptera: Formicidae) first arose 140-168 Ma (Moreau *et al*., 2006; Moreau & Bell, 2013) from solitary wasp ancestors (Johnson *et al*., 2013; Branstetter *et al*., 2017). At their origin, ants evolved eusociality (Evans, 1958; Lin & Michener, 1972; Hölldobler & Wilson, 1990; Danforth, 2002; Rehan & Toth, 2015), which is defined by a division of labor, cooperative brood care, and overlapping generations (Wilson, 1971). The transition to eusociality marks a significant evolutionary achievement (Szathmáry & Smith, 1995); only a handful of lineages have attained that level of social organization (Cameron, 1993; Chapman *et al*., 2000; Duffy, Morrison, & Ríos, 2000; Danforth, 2002; Brady *et al*., 2006; Inward, Beccaloni, & Eggleton, 2007). Eusociality has been crucial to the ecological success of the ants by allowing for an efficient division of labor (Macevicz & Oster, 1976; Oster & Wilson, 1978; Hölldobler & Wilson, 1990) and for emergent, complex behaviors at the level of the colony (Deneubourg & Goss, 1989; Gordon, 1992, 2002, 2007; Bonabeau, 1998; Page Jr & Mitchell, 1998; Theraulaz *et al*., 2002).

However, social living is not without costs, the largest of which is thought to be the increased burden of infectious disease, due to the increased frequency and density of potentially infectious contacts (Alexander, 1974; Freeland, 1976; Hoogland, 1979; Brown & Brown, 1986; Møller, Dufva, & Allander, 1993; Krause & Ruxton, 2002; Altizer *et al*., 2003; Patterson & Ruckstuhl, 2013; Schmid-Hempel, 2017). Extant ant genera are known hosts to a variety of parasitic organisms (Kistner, 1982; Schmid-Hempel, 1998), and their solitary ancestors were also likely host to parasites as well. As the ancestral ants transitioned to their increasingly social lifestyle it is reasonable to assume that this offered enhanced opportunities for disease transmission but also fostered the innovation of anti-parasite defenses, many of which were likely in place prior to the transition to eusociality (Meunier, 2015). Additionally, when the angiosperms arose in the Late cretaceous, ants rapidly diversified to occupy newly available ecological niches (Moreau *et al*., 2006). This diversification pushed ants into novel habitats and into contact with novel parasites.

Parasites have been increasingly recognized as important members of communities and ecosystems (Thomas, Adamo, & Moore, 2005; Hudson, Dobson, & Lafferty, 2006; Hatcher, Dick, & Dunn, 2012). Understanding drivers of parasite richness and community composition, as well as their cascading impacts, depends on understanding the ecological relationships they have with their hosts (Poulin, 1995). Furthermore, parasites are very important shapers of the evolutionary ecology of their hosts (Poulin, 2011; Schmid-Hempel, 2011). While parasites of many other social organisms (e.g. humans, lions, wolves), have been extensively studied, we lack a synthetic overview of the ant societies as hosts to a wide range of parasites.

An important question is whether the ants, one of the most ecologically successful groups of organisms on Earth, are lacunae of parasite biodiversity? Extensive taxonomic reports and natural history notes suggest that the diversity of parasites infecting ants is high (Kistner, 1982; Schmid-Hempel, 1998), but a comprehensive synthetic overview has been lacking. Thus, the aim of this work is to synthesize the known records of parasitic organisms infecting ants for both their taxonomic and functional biodiversity, in addition to reviewing the biology and life history of these organisms.

## 3. Methods

### 3.1 Defining parasite/parasitoid records

Many parasitic organisms and myrmecophiles associate with ants. Though these relationships can be quite nuanced (see SI), this work focuses only on parasitic organisms infecting individual ant hosts (cf. (Sherman, Seeley, & Reeve, 1988)). We define ‘parasite’ and ‘parasitoid’ following (Godfray, 1994; Lafferty & Kuris, 2002; Lafferty *et al*., 2015) A parasite is an organism that associates with one host during a given life stage, negatively impacts the host, and does not require host death as developmental necessity. In contrast, parasitoids also associate with one host during a given life stage but require host death in order to complete their development. While the parasite/parasitoid distinction is important, we will hereafter refer to all parasitic organisms generally as ‘parasites’ unless we are specifically discussing the differences between parasites and parasitoids.

We define a ‘record’ as a parasite/parasitoid genus (and species, if possible) associated with an ant genus (and species, if possible). For a given ant-parasite combination, multiple occurrences are possible, for example if the association is recorded from multiple locations, at multiple times, or by multiple collectors. However, we consider each ant-parasite combination as a singular ‘record' regardless of how many times and places that association has been observed. For interested readers, we provide references for each individual parasite-ant occurrence reported in the literature, but these are subsumed within a singular ant-parasite record (see SI).

### 3.2 Literature search

Records of parasites infecting ants were obtained by collecting all records within parasite-specific review papers and books (Poinar, 1975; Disney, 1994; Kathirithamby, 2009; Lachaud & Pérez-Lachaud, 2012) parasite-specific databases (the Universal Chalcidoidea database (Noyes, 2017), the Strepsiptera database (Kathirithamby, 2017), the Phorid catalog (Brown, 2018), and the USDA nematode collection (Handoo & Mowery, 2017)), host-specific books ((Kistner, 1982; Schmid-Hempel, 1998)), by executing a targeted search for all genera within the ants, and through serendipitous discovery (e.g. references in papers and books). The targeted search was carried out by systematically searching for each extant, recognized ant genus (Bolton, 2018)and the term par* within the CAB abstracts database (www.cabi.org), which provides access to English-language translation of paper abstracts which are in other languages and as such often not available in more commonly employed databases. In addition to both extant databases and the targeted ant genus search, serendipitous discoveries of ant-parasite occurrences were also made, often buried deep within older papers and books. It is likely that some records have inevitably been missed, but we are confident that the majority of published records have been included in this work. A schema for how the literature search was conducted is given in Fig. 1.

**Figure 1.**
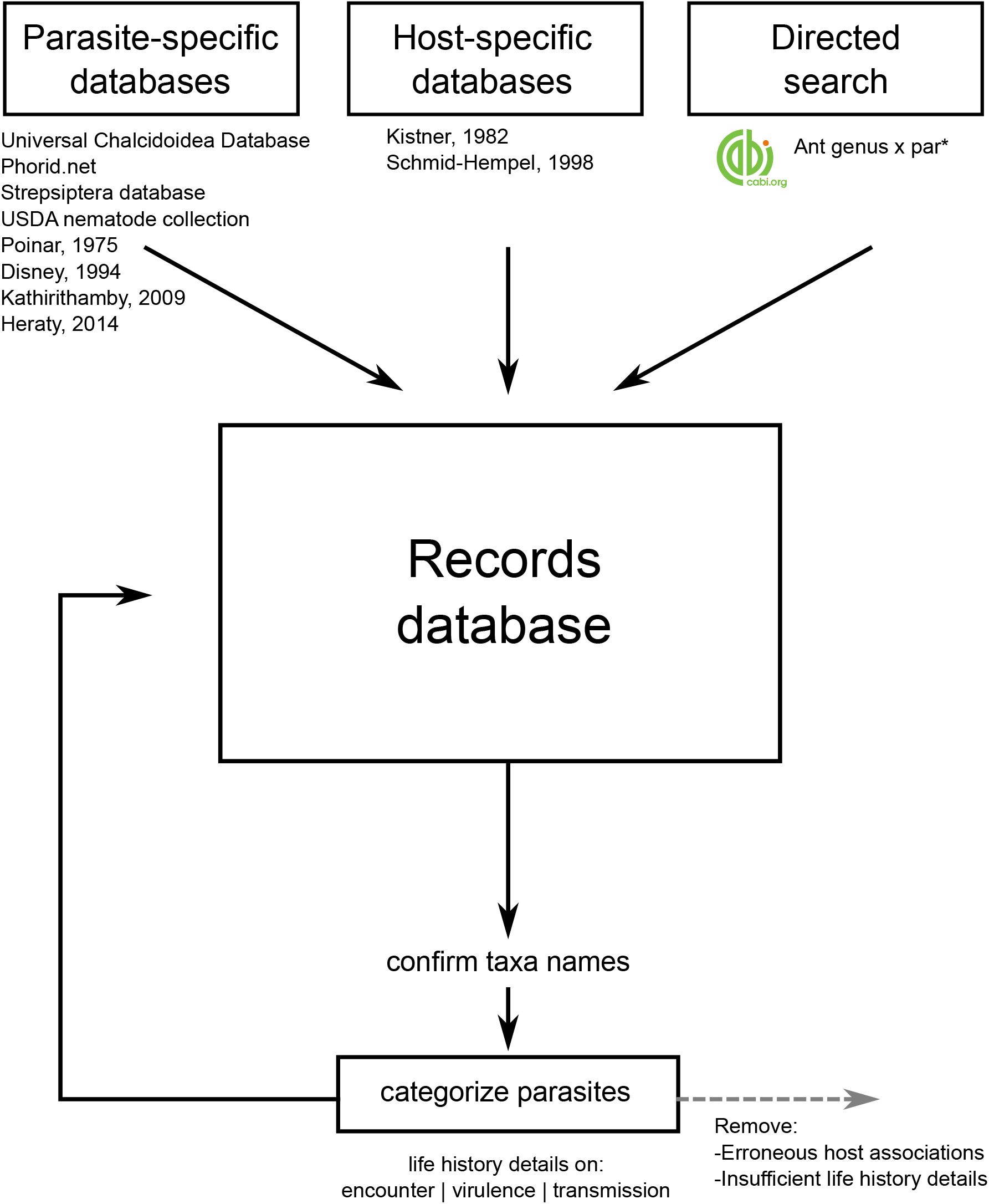
Literature search schema. Schema of how the literature search was conducted.

### 3.3 Categorizing parasite life history and epidemiological features

In order to synthesize trends in how parasites interact with their ant hosts, we scored relevant parasite life history traits. We focused on three essential epidemiological characteristics: encounter (how the parasite encounters hosts that it can infect, Fig. 2), virulence (morbidity or mortality for individual ants), and transmission (number of hosts in life cycle and whether the entire life cycle can occur within the confines of the ant nest, Fig. 3), discussed in the SI. Additionally, we report the ant life stage infected (i.e. brood vs. adult, worker vs. queen) if known, and categorize the relationship of the parasite and the ant according to the definitions previously discussed. The database of parasite records along with selected parasite traits is given in the SI.

**Figure 2.**
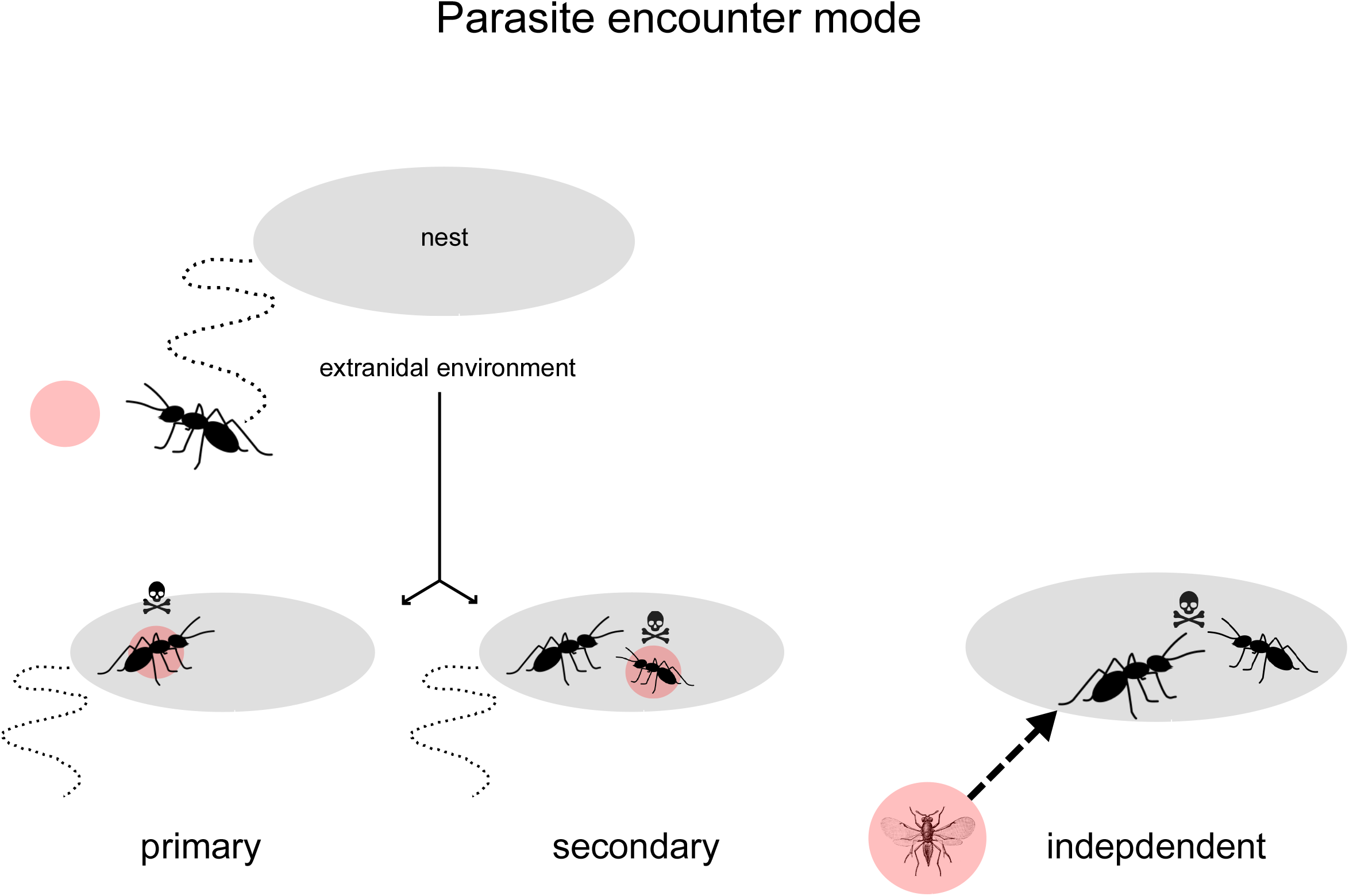
Parasite encounter modes. Schema of how parasites encounter their ant hosts (primary, secondary, independent). In primary encounters, the parasite encounters the host it will infect in the extranidal environment. In secondary encounters, the parasite encounters an individual in the extranidal environment, becomes phoretically attached to that individual, and is subsequently transferred to another ant host in the nest that it ultimately infects. In independent encounters, the parasite enters the nest independent of the actions of the host and infects its host once inside.

**Figure 3.**
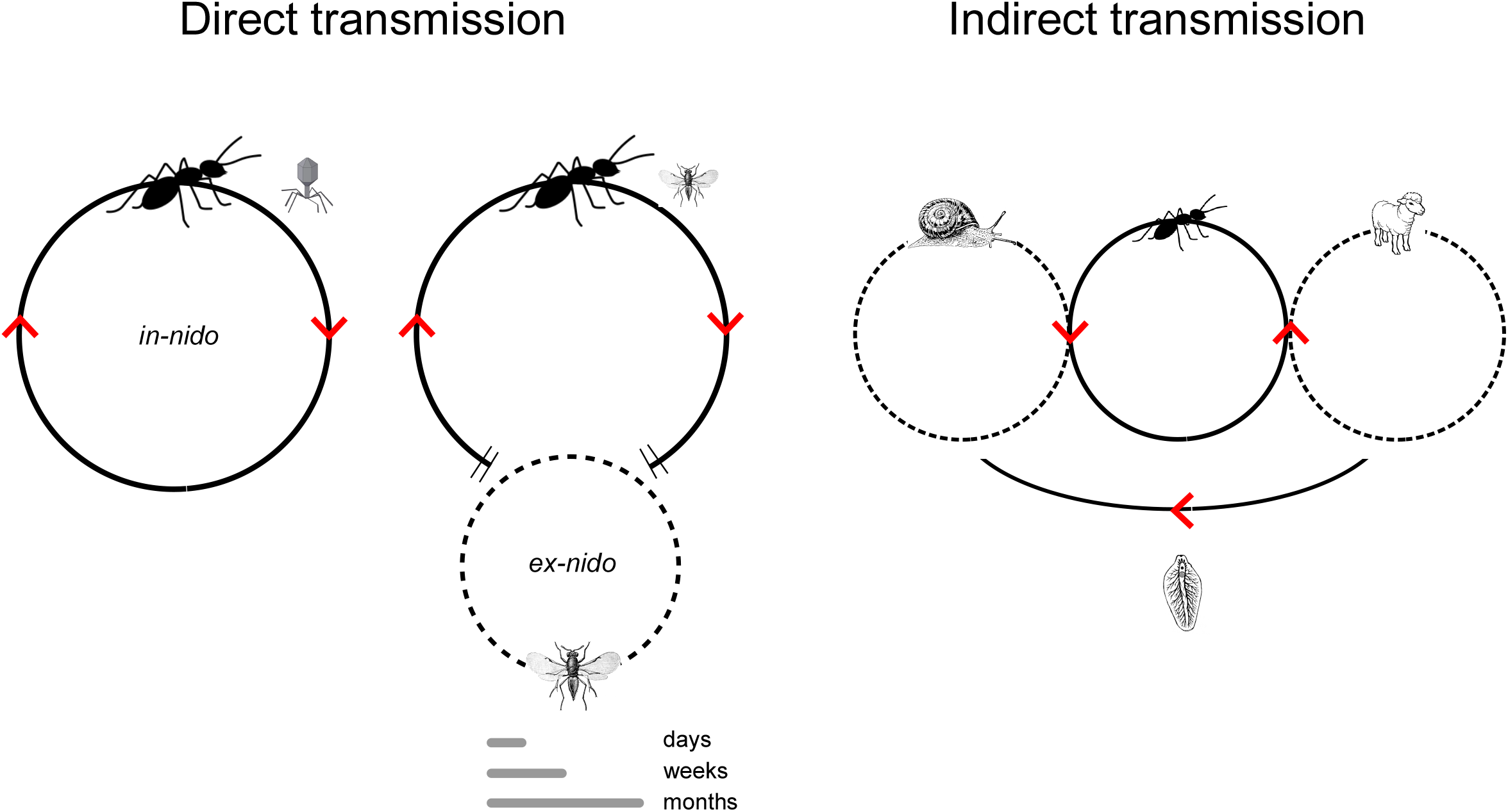
Parasite transmission mode and location. Schema of parasite transmission mode and location. Transmission can be described by how many hosts are needed to complete the parasite’s life cycle (direct transmission vs. indirect transmission) and by whether transmission to future hosts can take place within the confines of the nest (*in-nido* vs. *ex-nido* transmission). Parasites using direct transmission only require one host to complete their lifecycle whereas indirectly transmitting parasites require two or more hosts. For directly transmitting parasites, transmission to future hosts can take place either inside the nest (*in-nido* transmission), or if the parasite requires a period of time outside for either development or finding a mate, transmission may occur outside of the nest (*ex-nido* transmission). In the case of indirectly transmitting parasites, transmission to the next host must generally occur outside of the nest (*ex-nido* transmission), due to the multiple hosts used.

## 4. Results

We present our results in three major sections. We first provide host-specific results regarding the number of ant sub-families, genera, and species infected by parasites. Next, we present parasite-specific results, including trends in parasite traits and life history details, for each major group of parasites. Finally, we address bias in the reported parasite records and estimate potential biodiversity for parasites infecting ants.

### 4.1 Host-specific results

In this section, we take a host-centric view and look at how known parasite records assort across ant subfamilies, genera, and species. We report parasite records infecting 9 sub-families, 82 genera, and 580 ant species.

#### 4.1.1 Parasite records across ant subfamilies

Of the 17 currently recognized, extant ant subfamilies (Bolton, 2018), only 9 (52.94%) have any reported parasite, or parasitoid records (Table 1, Fig. 4). The subfamilies Formicinae and Myrmicinae had the highest number of records, with 523 and 520 records, respectively. These were followed by the Ponerinae (128 records) and and Dorylinae (86 records).

**Figure 4.**
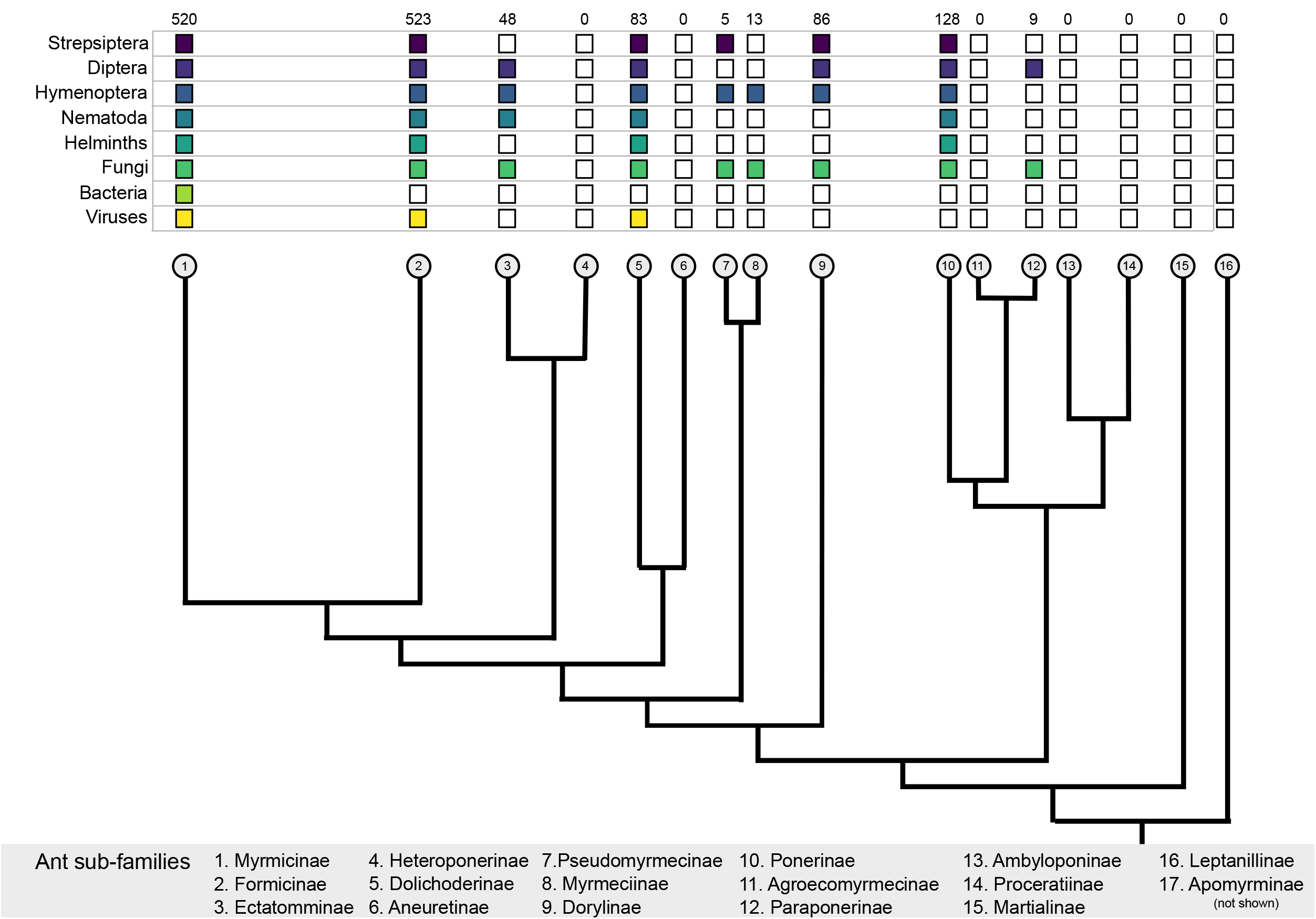
Parasite records by ant sub-family. The number and type of parasites recorded for each extant ant sub-family. The total number of records for each ant sub-family is listed at the top of the column; shaded boxes represent the presence of records for that particular parasite group infecting members of that ant sub-family. The fungi group encompasses the Fungi, Microsporidia, and Apicomplexa. The helminth group encompasses the Trematoda and Cestoda. The phylogeny in this figure was adapted from Moreau *et al*. 2006. *Science*.

**Table 1.** Parasite records by ant sub-family. A summary of the number of parasite records for each extant ant sub-family, and what percentage of total records each sub-family accounts for (total n. records = 1,415).

| **Sub-family** | **N. records** | **% of total records (n= 1,415)** |
| --- | --- | --- |
| Agroecomyrmecinae | 0 | 0.00% |
| Ambyloponinae | 0 | 0.00% |
| Aneuretinae | 0 | 0.00% |
| Apomyrminae | 0 | 0.00% |
| Dolichoderinae | 83 | 5.87% |
| Dorylinae | 86 | 6.08% |
| Ectatomminae | 48 | 3.39% |
| Formicinae | 523 | 36.96% |
| Heteroponerinae | 0 | 0.00% |
| Leptanillinae | 0 | 0.00% |
| Martialinae | 0 | 0.00% |
| Myrmeciinae | 13 | 0.92% |
| Myrmicinae | 520 | 36.75% |
| Paraponerinae | 9 | 0.64% |
| Ponerinae | 128 | 9.05% |
| Proceratiinae | 0 | 0.00% |
| Pseudomyrmecinae | 5 | 0.35% |
| **Total 1,415 records** |  | **100.00%** |

#### 4.1.2 Parasite records across ant genera

Of the 334 currently recognized extant ant genera (Bolton, 2018), we report only 82 (24.55%) as having parasites or parasitoids reported infecting them (Fig. S1, Table S1). Most of the ant genera with any reported parasite records had fewer than five records associated with them (median number of records = 5, Fig. S2). The ten ant genera with the most parasite records are *Camponotus* (192), *Solenopsis* (144), *Formica* (132), *Pheidole* (85), *Polyrhachis* (69), *Lasius* (67), *Atta* (61), *Pachycondyla* (60), *Acromyrmex* (59), and *Odontomachus* (32).

#### 4.1.3 Parasite records across ant species

While there are currently over 15,000 recognized extant ant species (Bolton, 2018), we only report parasite records associated with 580 ant species. Thus, fewer than 5% of the total estimated ant species have any recorded parasites or parasitoids.

### 4.2 Parasite-specific results

We found 1,415 records of parasitic organisms infecting ants, comprising 51 families, 159 genera, and 615 species infecting 82 ant genera and 580 ant species. In Fig. 5, we plot the number of records associated with each major parasite group. In Table 2 we summarize the number of parasite records and the unique ant genera and species associated with each parasite family. A compendium of all records and their relevant life history traits is given in Table S2.

**Figure 5.**
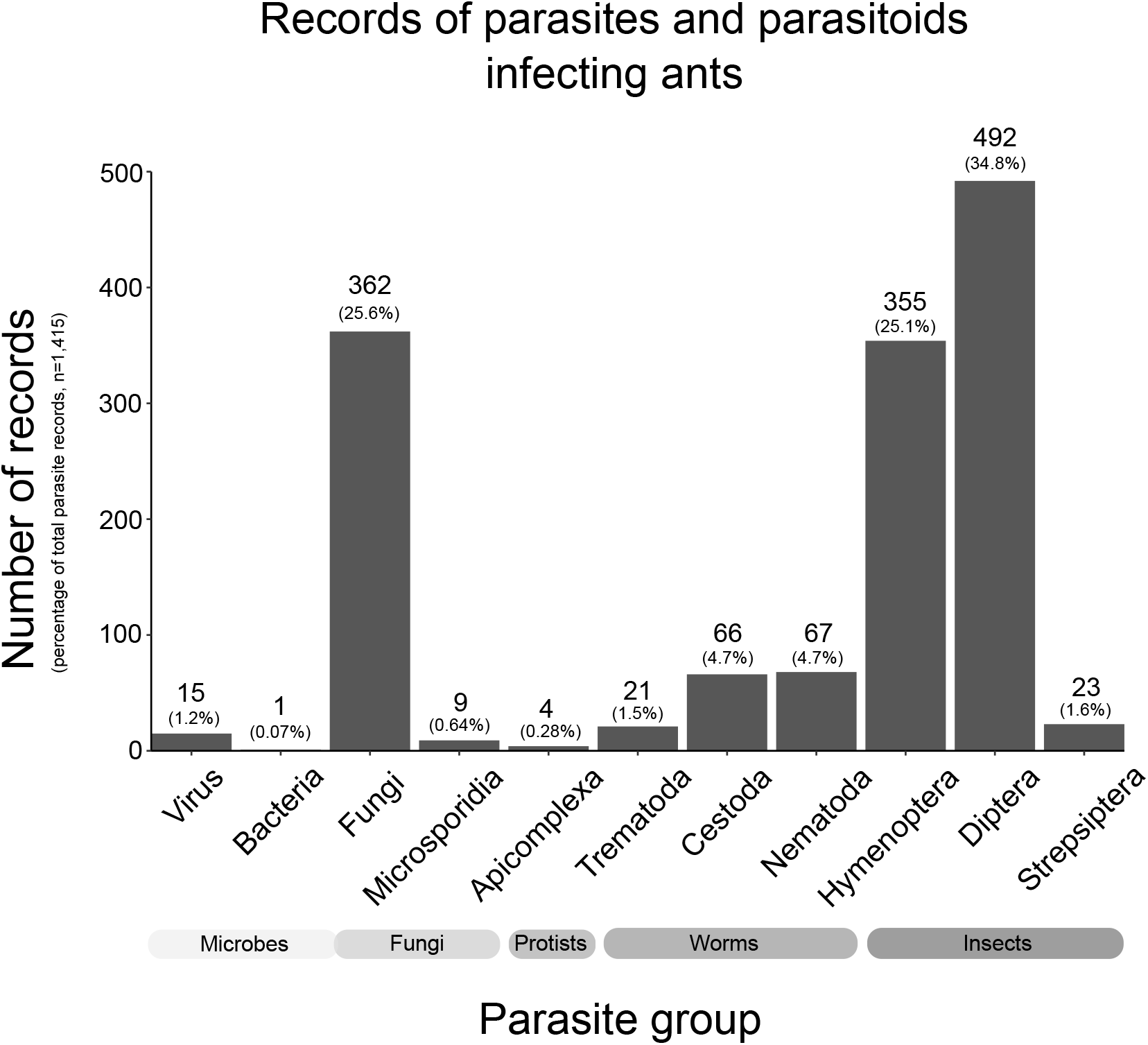
Parasite records for each major parasite group. The number of parasite records (and percentage of total records, n=1,415) for each major parasite group. Prot. = protist.

**Table 2.**
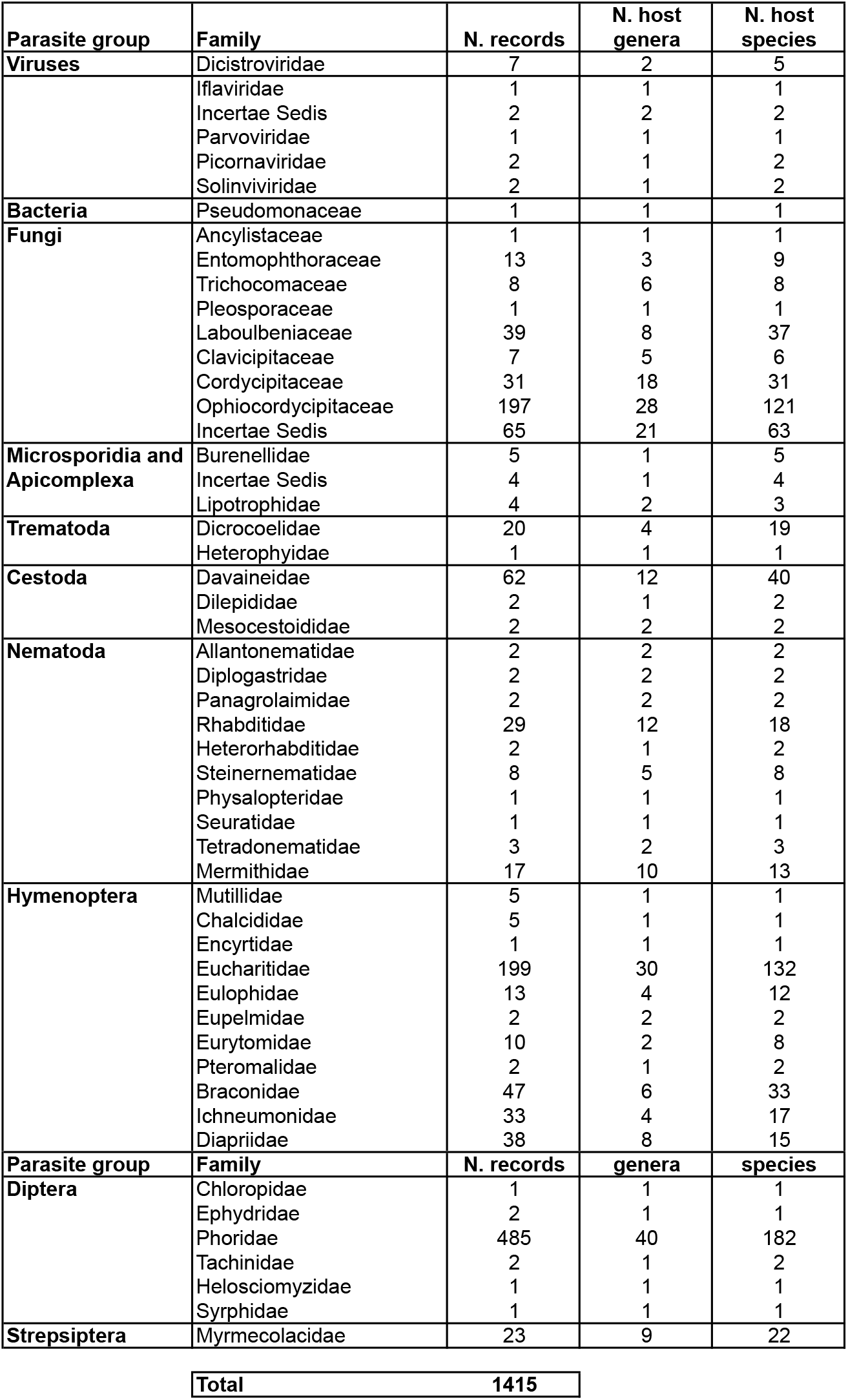
Parasite records by parasite family. A summary of the number of records, unique host genera infected, and unique host species infected for each parasite family.

#### 4.2.1 overall parasite life history trends

Below we summarize general trends in life history traits of parasites infecting ants. We summarize the percentage of parasite records with a given life history trait (e.g. direct transmission) in Table 3. In Figs. 6 – 10 we show how these traits assort across the major parasite groups.

**Figure 6.**
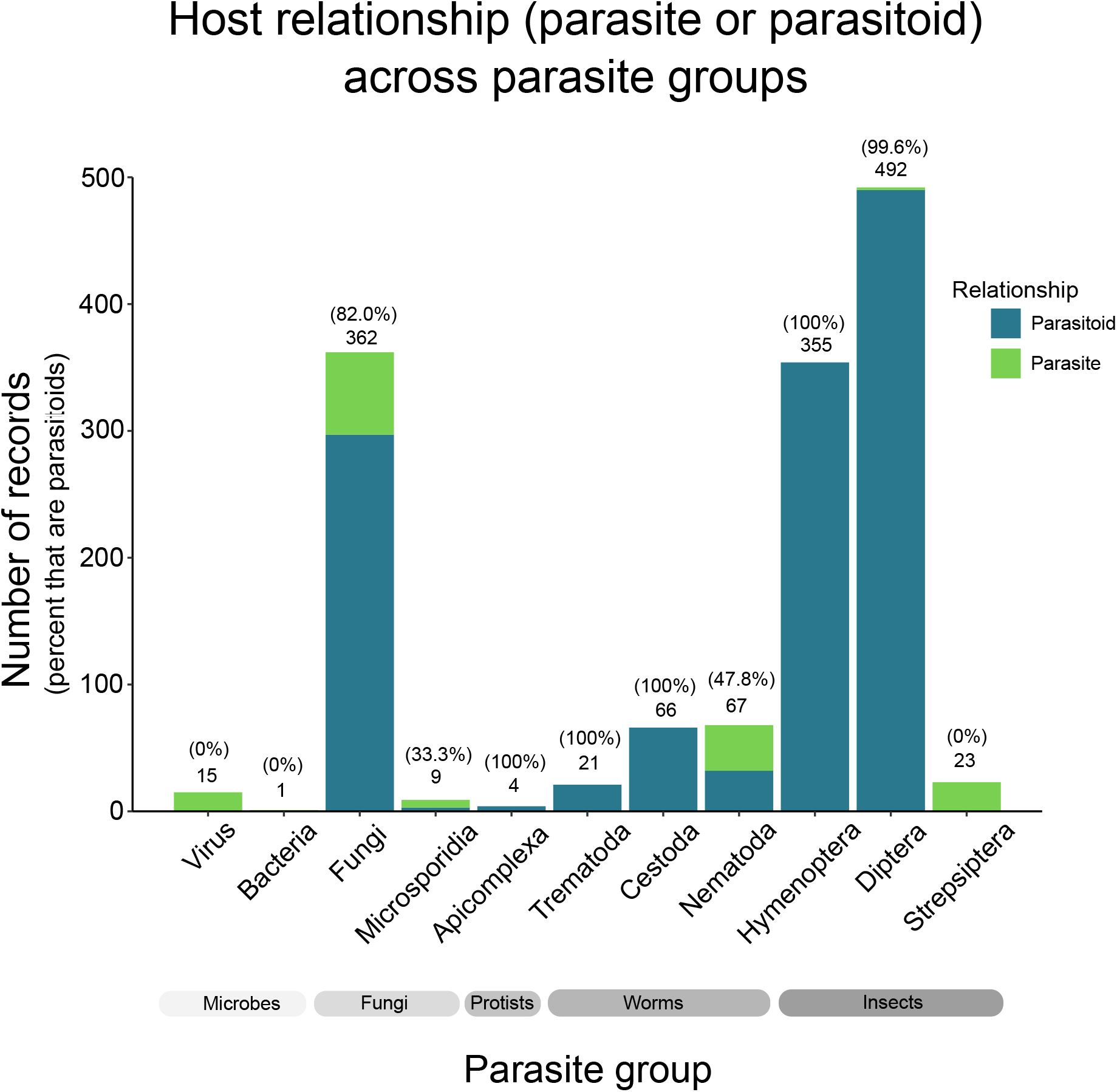
Host relationship by parasite group. The number of parasite records for each major parasite group, colored by relationship to the host (parasite vs. parasitoid). The number at the top of each bar is the total number of records for that parasite group; the percentage in parentheses represents the percentage of those parasite group’s records which are scored as parasitoids.

**Figure 7.**
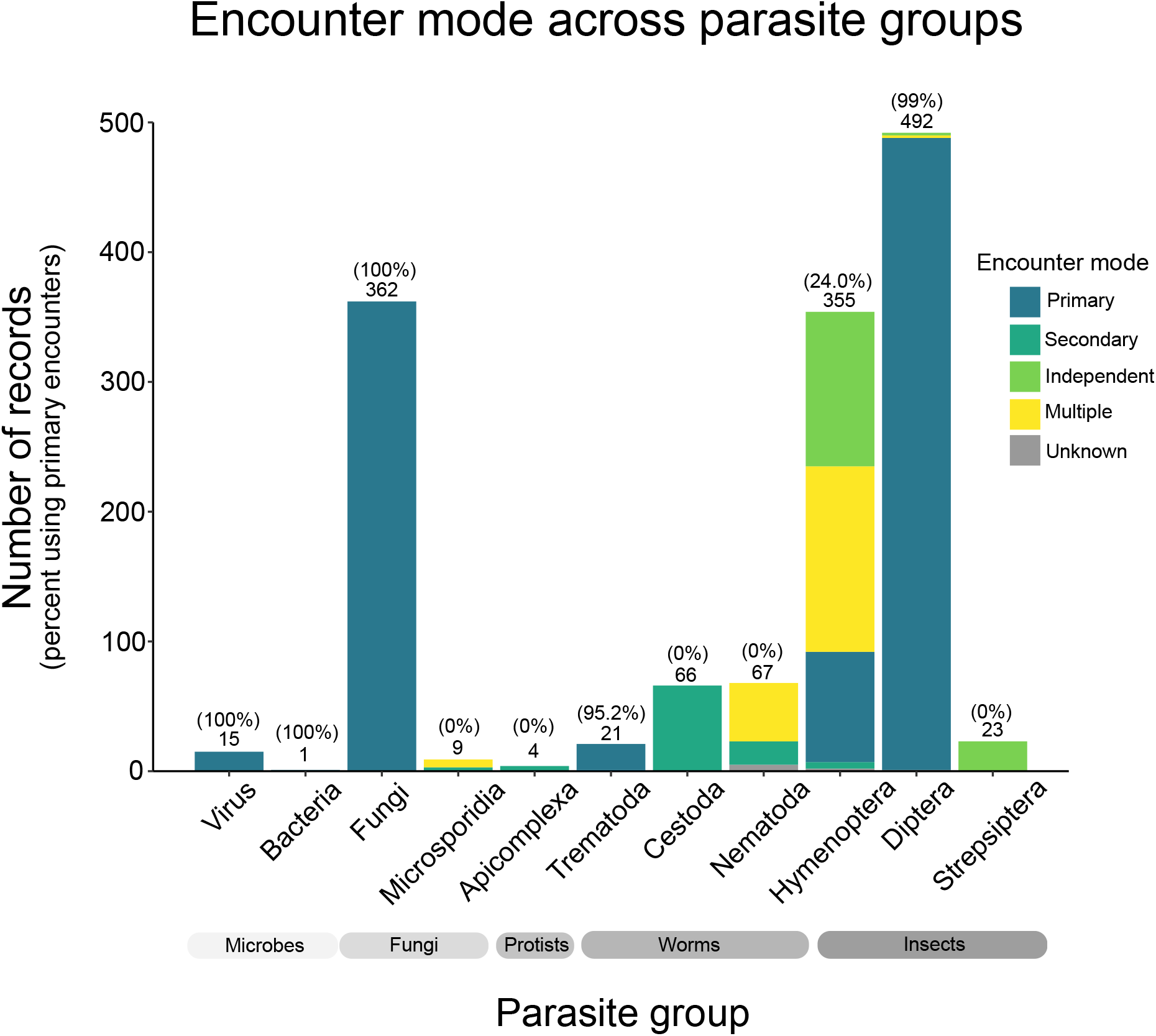
Host encounter mode by parasite group. The number of parasite records for each major parasite group, colored by host encounter mode (primary, secondary, independent, multiple, unknown). The number at the top of each bar is the total number of records for that parasite group; the percentage in parentheses represents the percentage of those parasite group’s records which are scored as primary encounters.

**Figure 8.**
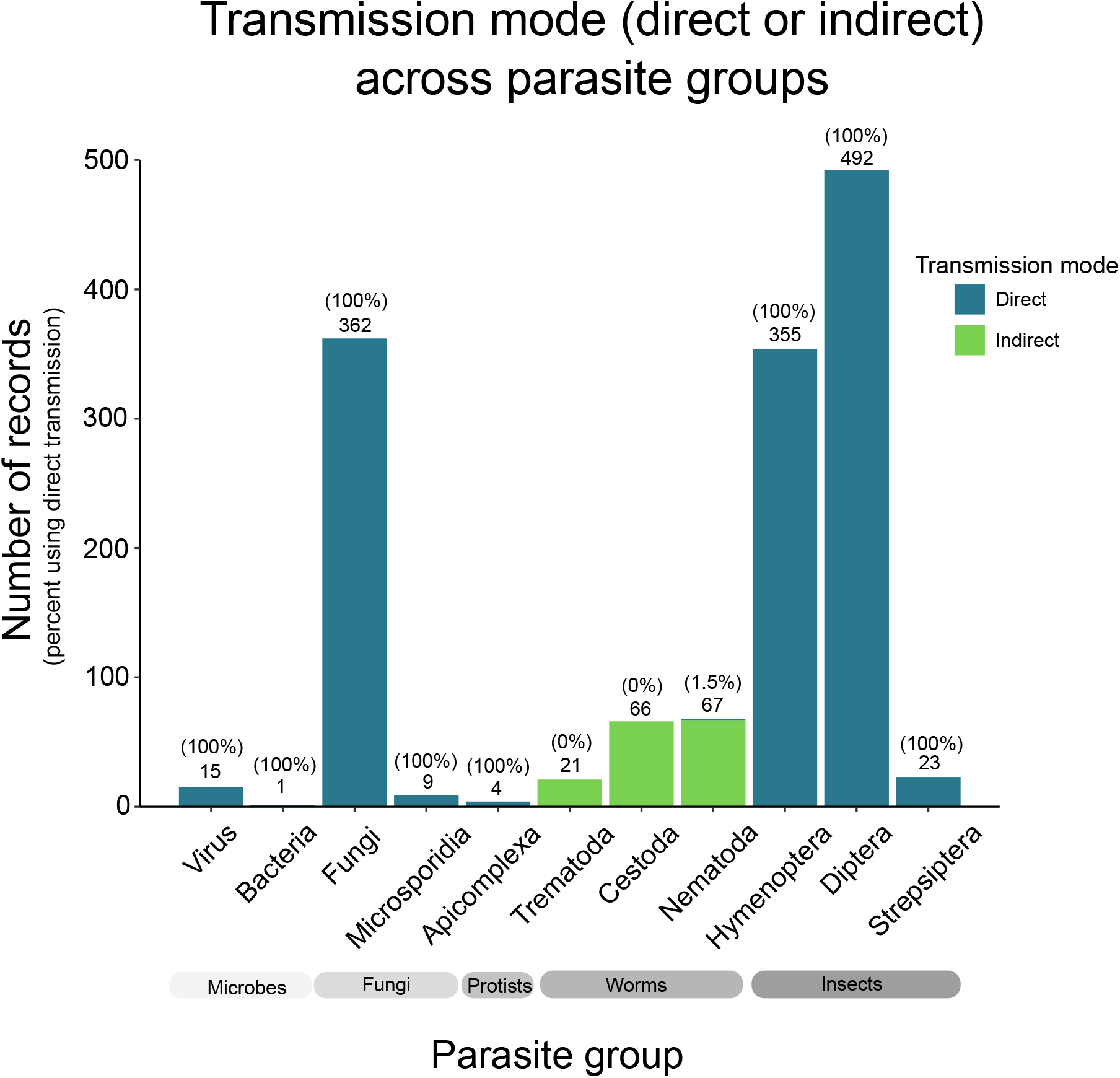
Parasite transmission mode by parasite group. The number of parasite records for each major parasite group, colored by parasite transmission mode (direct vs. indirect). The number at the top of each bar is the total number of records for that parasite group; the percentage in parentheses represents the percentage of those parasite group’s records which are scored as directly transmitting.

**Figure 9.**
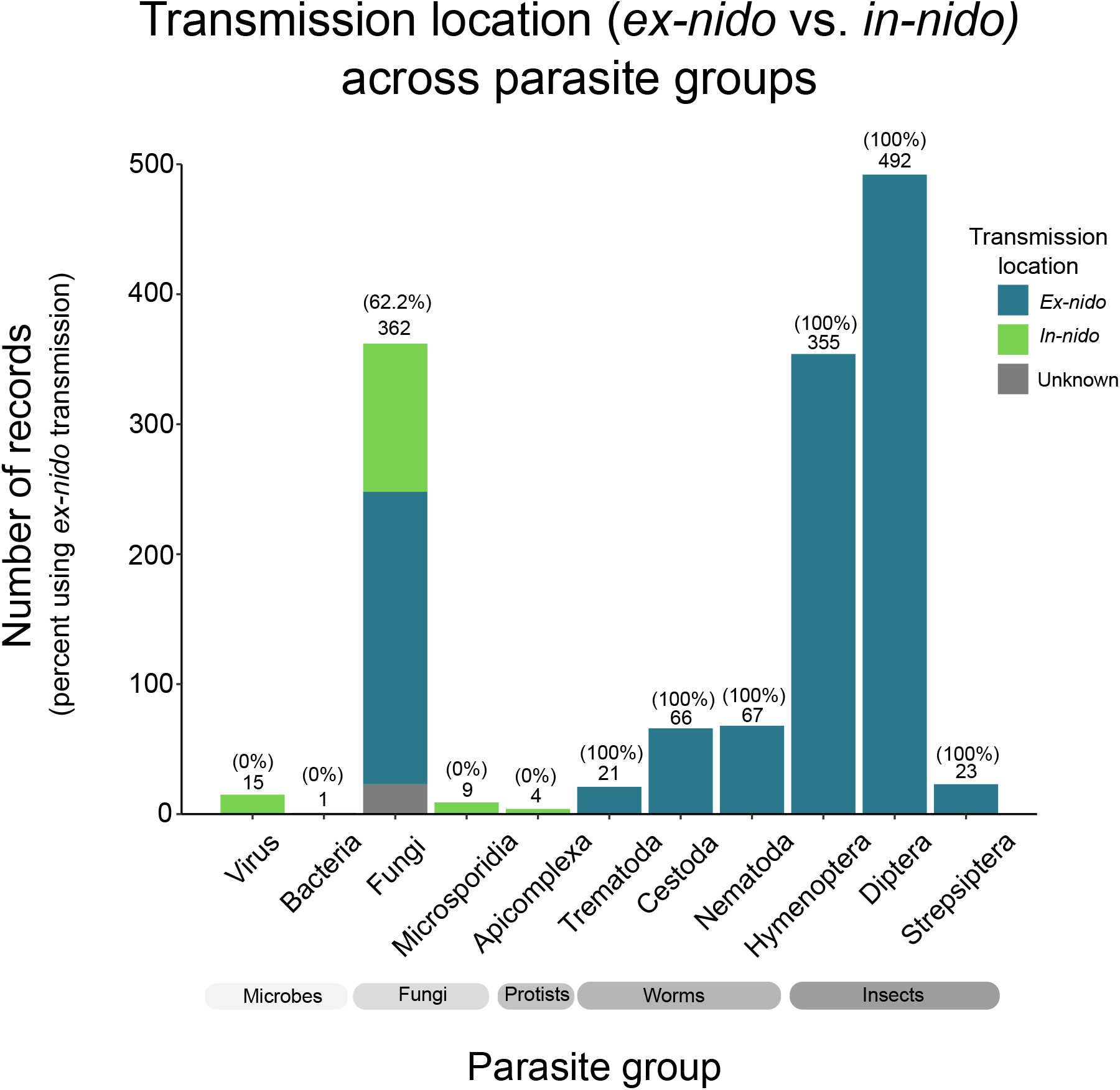
Parasite transmission location by parasite group. The number of parasite records for each major parasite group, colored by parasite transmission location (*in-nido* vs. *ex-nido* vs. unknown). The number at the top of each bar is the total number of records for that parasite group; the percentage in parentheses represents the percentage of those parasite group’s records which are scored as using *ex-nido* transmission.

**Figure 10.**
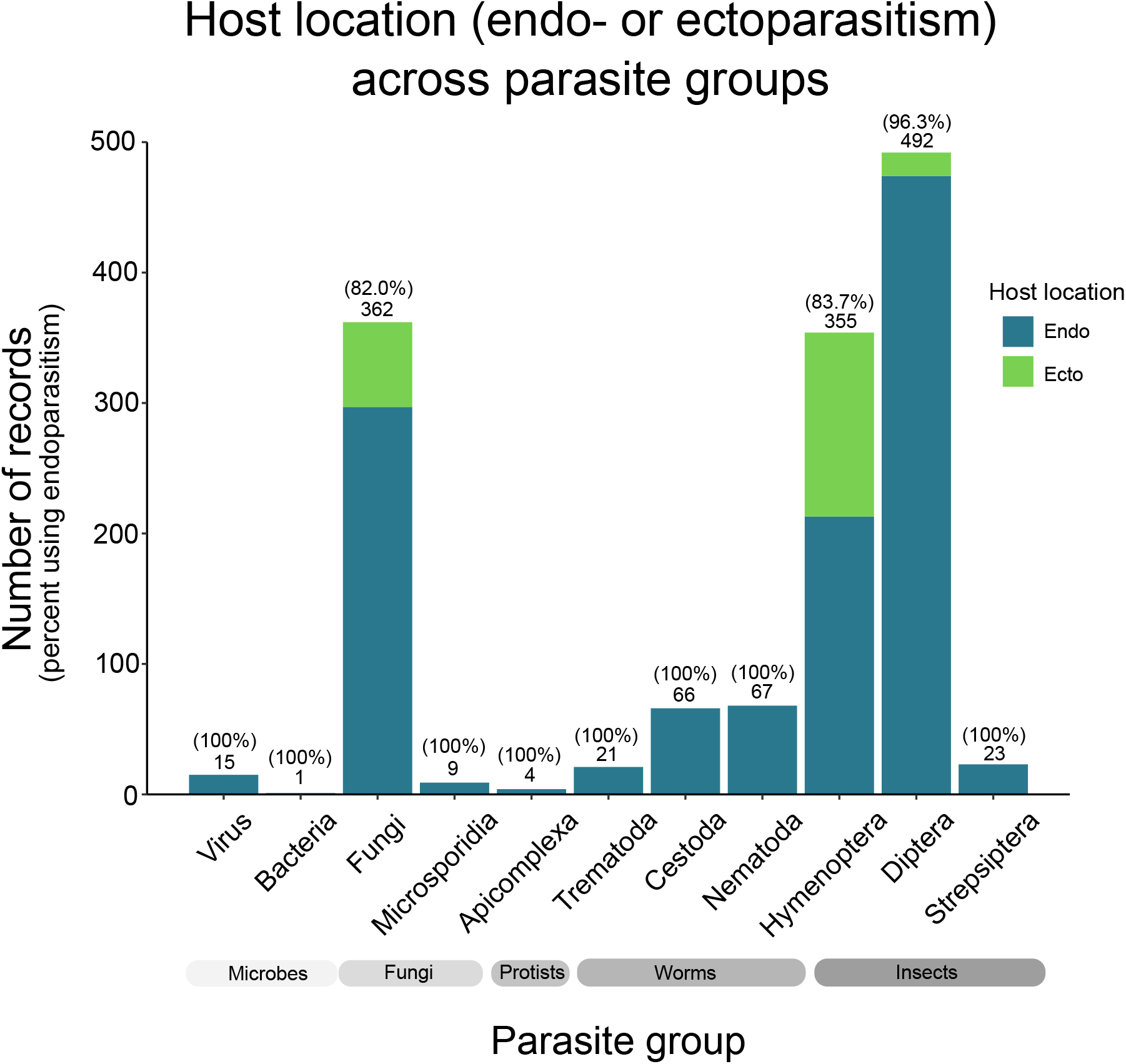
Endo-vs. ectoparasitism by parasite group. The number of parasite records for each major parasite group, colored by parasite location in or on the host (endo-vs. ectoparasitism). The number at the top of each bar is the total number of records for that parasite group; the percentage in parentheses represents the percentage of those parasite group’s records which are scored as being endoparasites.

**Table 3.**
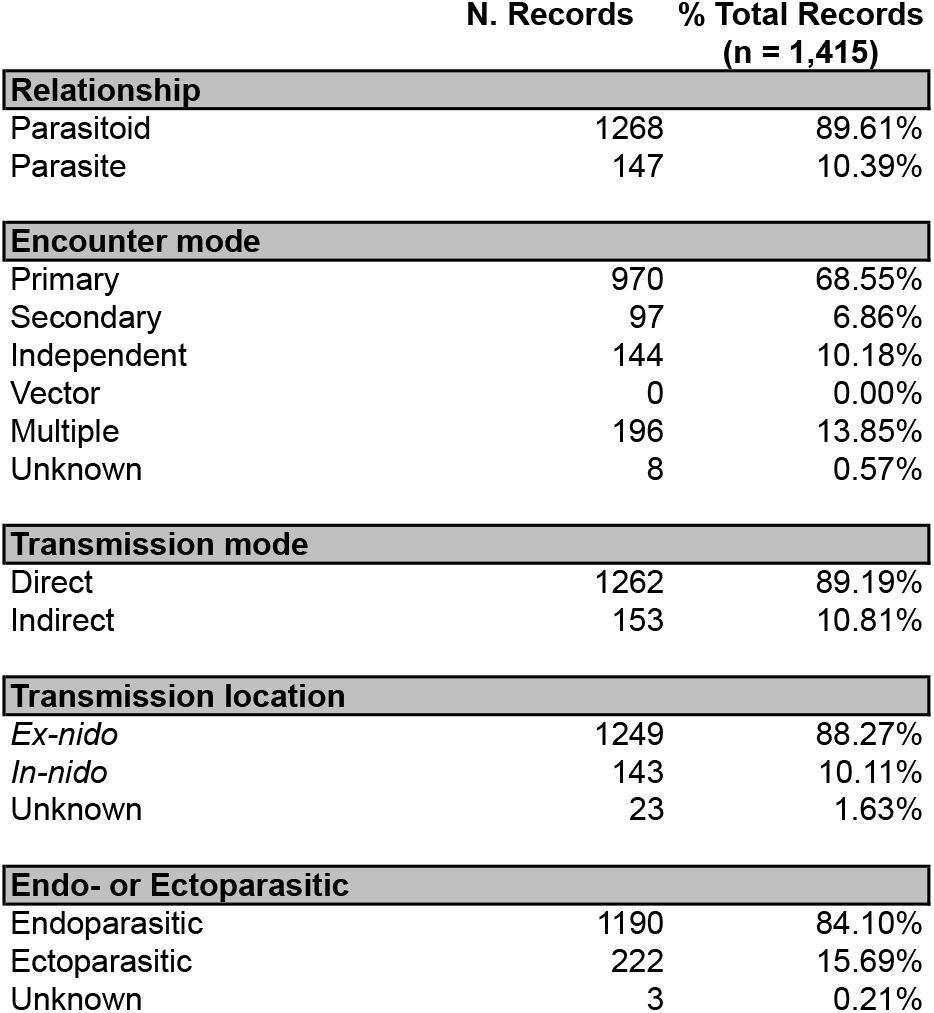
Parasite life history traits. A summary of the number of records scored with each parasite life history/epidemiological trait, and what percentage of the total records each of those traits comprises (total n. records = 1,415).

Parasitoidism, in which the death of the ant host is required as a developmental necessity, predominates, accounting for 89.61% of reported records (1,268/1,415) (Fig. 6). In contrast, parasitism accounts for only 10.39% of the reported records (147/1,415). In order to infect ant hosts, the majority of parasitic organisms use primary encounters (68.55%, 970/1,415 records, Fig. 7), wherein ants encounter parasitic organisms in the extranidal environment and subsequently become infected. 6.86% use secondary encounters (97/1,415 records), in which parasitic organisms encounter ants in the extranidal environment, become phoretically attached, and are carried inside the nest where they subsequently infect other individuals. 10.18% use independent encounters (144/1,415 records), meaning that the parasitic organism actively enters the ant nest on their own. 13.85% use multiple means of encountering their hosts (196/1,415 records), and for 0.57% of parasite records it remains unknown how they encounter their hosts (8/1,415 records).

Most parasitic organisms infecting ants use direct transmission (only a single host is needed to complete life cycle), comprising 89.19% of the reported records (1,262/1,415 records, Fig. 8). In contrast, a smaller proportion use indirect (multi-host) life cycles, and these parasites are found within the Trematoda, Cestoda, and Nematoda (10.81%, 153/1,415, Fig. 8). Though most parasites infecting ants only require a single host to complete their life cycle, the vast majority of parasites use *ex-nido* transmission, which requires leaving the nest for a period of time to develop or mate before transmission to next ant host can occur (88.27%, 1,249/1,415 records, Fig. 9). *Ex-nido* transmission is found within the fungi, worms, and insects, but is absent from viruses, bacteria, microsporidia, and apicomplexa (Fig. 9). Future work could determine if *ex-nido* transmission occurs in these groups. Finally, the majority of parasites and parasitoids infecting ants develop internally inside the host as endoparasites (84.10%, 1,190/1,415 records, Fig. 10).

#### 4.2.2 Host specificity

Understanding whether a parasite infects a single genus or species or many host genera and species is important for estimating the potential biodiversity of parasites infecting ants. The majority of parasites infecting ants appear to be specialists, with each parasite genus typically only infecting one or a few host genera (median host genera infected = 1, mean = 2.72, Fig. 11a), and infecting few host species (median host species infected = 2, mean = 6.42, Fig. 11b). However, there are some parasites that appear to be more cosmopolitan with their host choice (Fig. 12a,b). For example, some fungal and dipteran genera have been recorded infecting over 15 different ant genera (Fig. 12a). However, based on the parasite records thus collected, most ant-infecting parasite genera seem to infect only a few ant genera and/or species.

**Figure 11.**
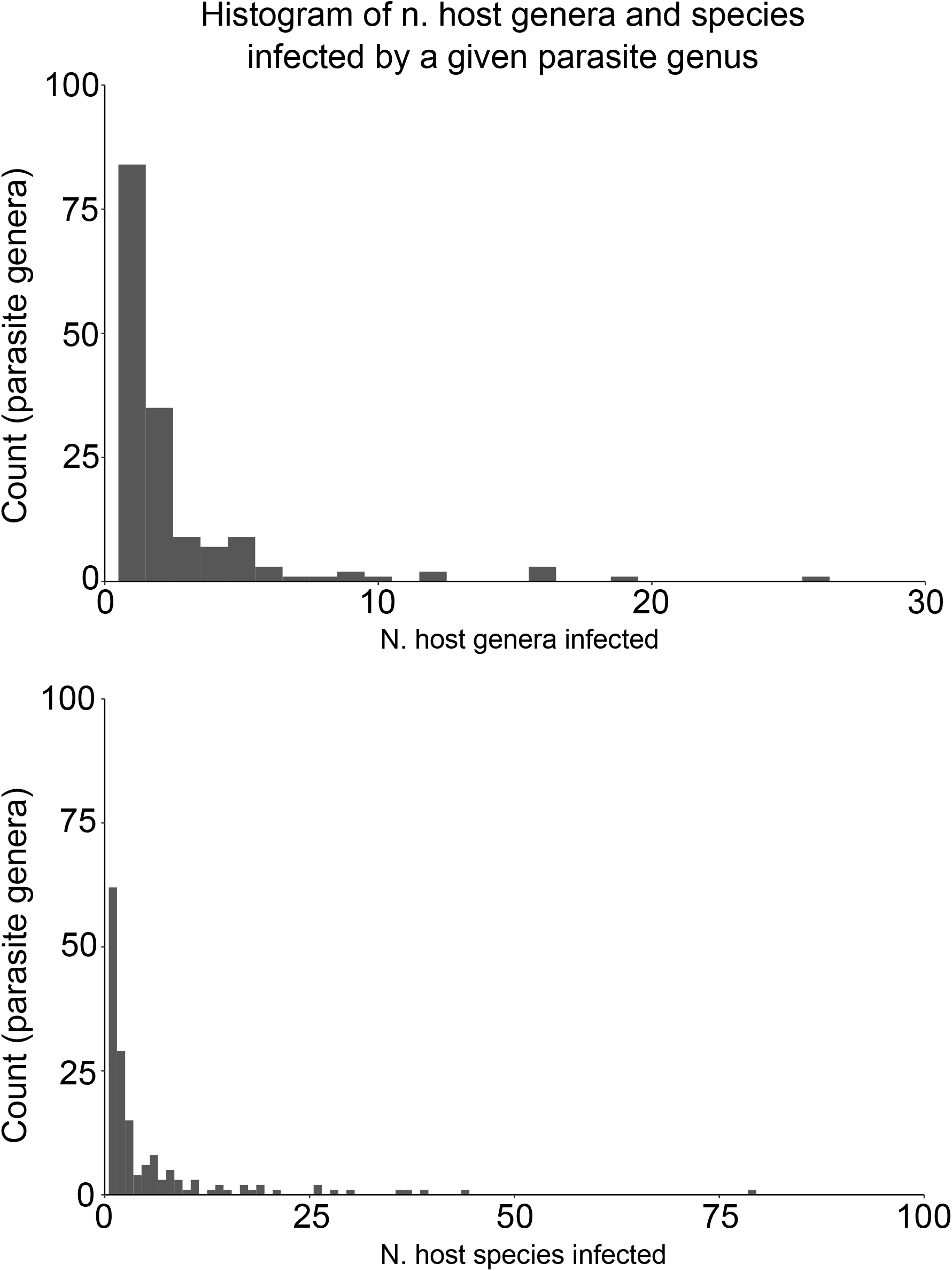
Histogram of n. host genera and species infected by a given parasite genus. Histogram of the number of (a) host genera and (b) host species infected by a given parasite genus. Most parasite genera infect a small number of host genera (mean = 2.72 host genera, median = 1 host genera). Most parasite genera infect a small number of host species (mean = 6.42 host species, median = 2 host species).

**Figure 12.**
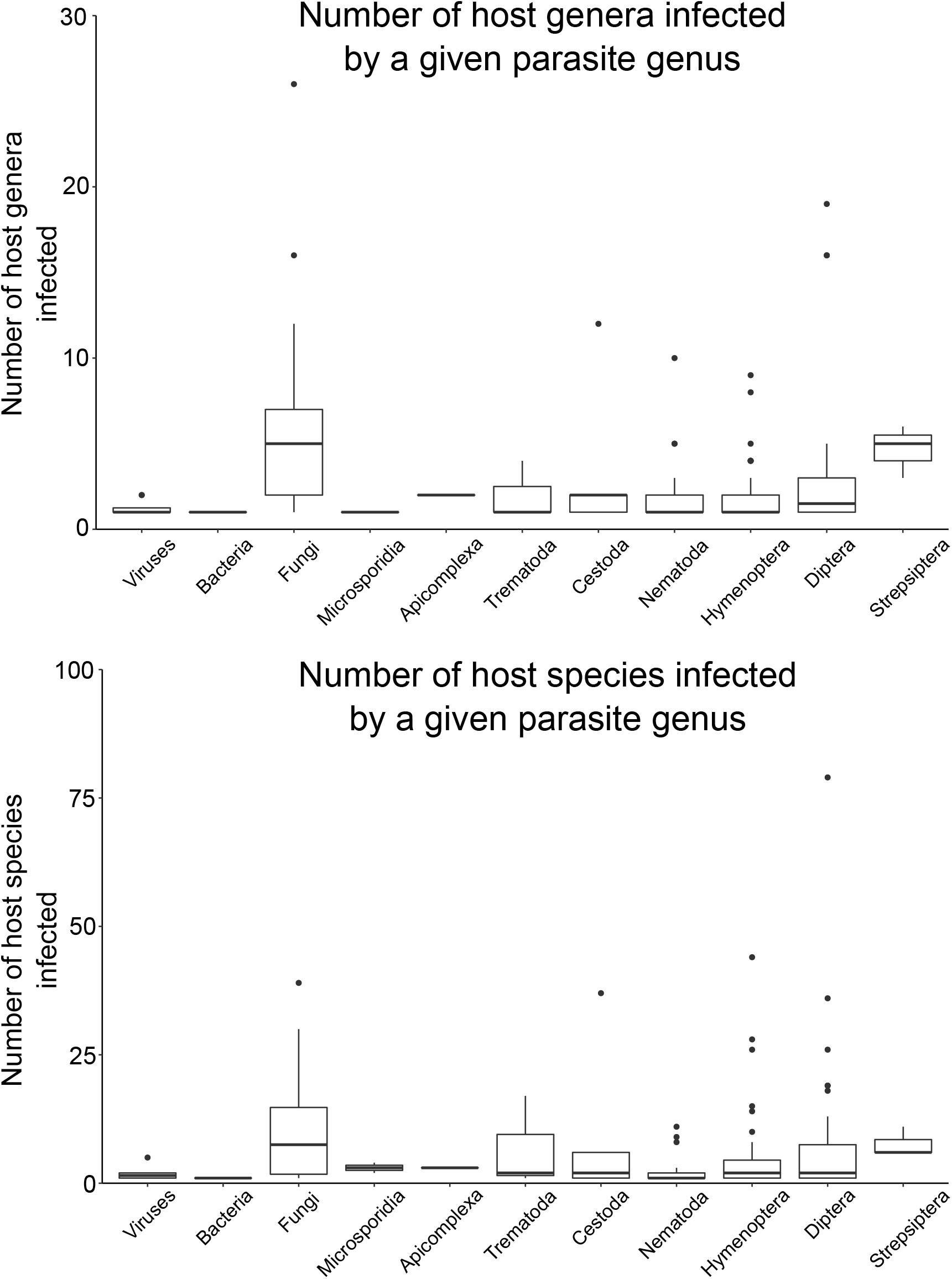
Boxplots of host genera and species infected by a given parasite genus, by parasite group. Boxplots of the number of (a) host genera and (b) host species infected by a given parasite genus, for each major parasite group. Black lines represent median values, boxes represent the range of values in the 1st through 3rd quartiles, and whiskers represent 1.5 times the inter-quartile range.

### 4.3 Viruses

Knowledge of viruses infecting ants is quite limited, largely due to the relative difficulty in isolating and identifying viruses compared to other taxa and due to the inherent difficulties in observing and sampling dead or dying ant hosts, particularly in natural field settings. At least 18 viruses are known to infect genera within the Apidae (Bailey, 1976; Allen & Ball, 1996; Schmid-Hempel, 1998; Ellis & Munn, 2005; Chen & Siede, 2007), but our knowledge of viruses potentially infecting the Formicidae is just beginning to catch up. The first reports of potential viral infections in ants were by Steiger et al. (Steiger *et al*., 1969), who found ‘virus-like' particles infecting *Formica lugubris* and Avery et al. (Avery *et al*., 1977), who found `virus-like' particles infecting an unidentified *Solenopsis* species, but the first confirmed report did not occur until 2004.

#### 4.3.1 Single-strand RNA viruses (ssRNA+)

Valles et al. (Valles *et al*., 2004) report the first confirmed instance of a virus infecting ants, *Solenopsis invicta* virus-1 (SINV-1). Subsequent work has found SINV-1 infecting S. *invicta* colonies in both Argentina and the United States (Valles *et al*., 2007, 2009). The virus has been found infecting other species within the *Solenopsis* genus (*S. geminata, S. richteri*, and S. *carolinensis*, as well as some *Solenopsis* species hybrids, (Valles, 2012). While SINV-1 has been detected in ants, its actual host pathology remains unclear. Valles et al. (Valles *et al*., 2004) note no observable symptoms of virus infection in natural populations, but did note virus-induced larval mortality in the lab. While the virus appears to have a limited impact on individual hosts, it does appear that its presence may alter the competitive ability of infected colonies. When competed against colonies of other species, SINV-1 infected colonies of S. *invicta* were weaker competitors compared to their uninfected counterparts (Chen, Kafle, & Shih, 2011).

Shortly after the discovery of SINV-1, two other related viruses, SINV-2 and SINV-3, were found to infect *Solenopsis invicta* (Valles *et al*., 2007; Valles & Hashimoto, 2009). There is little evidence that SINV-2 causes any observable host pathology and host specificity remains unknown (Valles, 2012). Allen and colleagues (Allen, Valles, & Strong, 2011) report finding co-infections of various combinations of SINV-1, SINV-2, and SINV-3 in individuals of *Solenopsis invicta* colonies, with polygyne colonies having a higher virus prevalence. It is unknown how having multiple infections impacts host morbidity and mortality, or how the presence of multiple viruses affects colony-level functioning.

In contrast, SINV-3 has been shown to cause considerable mortality in laboratory colonies of S. *invicta*, particularly among brood (Valles & Hashimoto, 2009). Further studies have shown that SINV-3 can negatively impact queen egg production and brood development (Valles *et al*., 2013a), perhaps due to behavioral changes caused by the virus that impede the ability of workers to disseminate food within the colony (Valles, Porter, & Firth, 2014). In some cases, SINV-3 infected colonies have been shown to rebound following the initial infection period (Valles, 2012), but SINV-3 remains the most pathogenic virus known to infect ants.

Celle et al. (Celle *et al*., 2008) report Chronic Bee Paralysis virus (CBPV) from *Camponotus vagus* and *Formica rufa*. This virus was isolated from ants due to their proximity to infected beehives and not because there were signs of virus-induced pathology to the ants. The effects of CBPV on ant health and the transmission potential inside ant nests remain unknown.

Gruber et al. (Gruber *et al*., 2017) conducted an extensive survey for viruses of the invasive Argentine ant *Linepithema humile* in New Zealand. They found a novel ant-specific virus *Linepithema humile* virus-1, as well as several viruses normally found in bees: Kashmir bee virus (KBV), black queen cell virus (BQCV), and deformed wing virus (DWV). The pathology of these viruses on their ant hosts is not known.

#### 4.3.2 Single-strand DNA viruses (ssDNA +/−)

Valles et al. (Valles *et al*., 2013b) report the first DNA virus from a hymenopteran host, *Solenopsis invicta* densovirus (SíDNV) infecting S. *invicta* in Argentina. Despite extensive sampling of S. *invicta* colonies in the United States where the ant is invasive, the virus has not been detected there, suggesting that the virus might be a promising tool for biocontrol of the pestiferous ant species. It is unclear to what extent the virus impacts individual ant morbidity or mortality.

#### 4.3.3 Records of viruses infecting ants [15 records]

We report 15 records of viruses infecting ants, representing 9 unique viruses (SINV-1, SINV-2, SINV-3, SiDNV, LHUV-1, KBV, DWV, BQCV, and CBPV, Table S2). There is a wide range in the reported pathogenicity of these viruses, ranging from no observable symptoms to host mortality. Thus far, only 7 ant species (and 2 hybrid *Solenopsis* species) have been identified as hosts. This is certainly an underestimate of the number of viruses infecting ants and an underestimate of the number of ant species infected; numbers will grow as more surveys explicitly looking for viral pathogens are performed.

### 4.4 Bacteria

Reports of bacteria associated with ants have become more prevalent over the past few decades, as improved molecular methods have allowed for the identification of bacteria species previously unculturable or unidentifiable in the lab. Several surveys have looked for bacterial pathogens and associates within ant colonies (Jouvenaz, Banks, & Atwood, 1980; Baird, Woolfolk, & Watson, 2007; Ishak *et al*., 2011; Powell, Hanson, & Bextine, 2014; Woolfolk *et al*., 2016b), often for purposes of identifying potential biocontrol agents. However, in the majority of these surveys, the exact association between the bacteria and the ant host (i.e. mutualistic, pathogenic, or somewhere in between) remains unknown.

Lofgren et al. (Lofgren, Banks, & Glancey, 1975) report that a *Pseudomonas* bacteria was found in dead ants taken from *Solenopsis* colonies in Mississippi, USA. When cultured and fed to ants in the lab, the bacterium readily killed all larvae and 50% of the adult workers. However, it is unclear whether this bacterium was the causative agent of death in the dead ants they initially isolated from the nest mounds, or whether *Pseudomonas* colonized these dying/dead ants opportunistically. While mortality was observed in the lab, it is unclear whether the infective dose used represents a biologically reasonable dose. Jouvenaz et al. (Jouvenaz *et al*., 1980) conducted a wide survey of bacterial and microsporidian pathogens associated with *Solenopsis invicta* and *S. richteri* in their native range in Brazil. Out of 640 colonies sampled in their surveys, a few larvae in only one colony were found to harbor a ‘sporeforming bacterium' (Jouvenaz *et al*., 1980). Attempts to isolate the bacterium were unsuccessful, so its transmission and pathogenicity could not be determined.

More recently, surveys looking for ant microbial associates have identified many bacteria in the environment in and around ant nests. Baird et al. (Baird *et al*., 2007) found 58 bacterial isolates associated with *Solenopsis* mound soil, mound plant debris, and on the external body regions of ants. Ishak et al., Powell et al., and Woolfolk et al. (Ishak *et al*., 2011; Powell *et al*., 2014; Woolfolk *et al*., 2016b), performed follow-up studies, finding bacterial isolates from *Solenopsis invicta* and *Solenopsis geminata* mound soil, surrounding plant debris, and internal and external ant tissues. Ongoing work is examining *Wolbachia* in ants, but as these are peculiar reproductive parasites that are vertically transmitted, we do not consider them here.

#### 4.4.1 Bacteria as endosymbionts and mutualists

Much work is currently being devoted to understanding the diversity of bacteria within ant colonies and the role that they provide as endosymbionts and mutualists (Van Borm, Billen, & Boomsma, 2002; Wenseleers, Sundström, & Billen, 2002; Hughes, Pierce, & Boomsma, 2008; Russell *et al*., 2009; Ishak *et al*., 2011; Kautz *et al*., 2013). Obligate endosymbionts, such as *Blochmannia* (Sauer *et al*., 2000) and *Wolbachia* (Wenseleers *et al*., 2002), have long co-evolutionary histories with their ant hosts and have been linked to the evolution of herbivory in the ants (Russell *et al*., 2009), and may be extremely important for the nutritional ecology of different ant species (Feldhaar, 2011). Other bacteria species may play an important role in preventing colony infection by other pathogens and parasites (Little *et al*., 2006; Mattoso, Moreira, & Samuels, 2012). How ants and other social insects differentiate between their endosymbiotic bacteria and those that are pathogenic is an interesting and open question (Feldhaar & Gross, 2008), as is how these relationships evolve over time (Hughes *et al*., 2008).

#### 4.4.2 Records of bacteria infecting ants [1 record]

We report only a single case where a bacterium is known to cause host morbidity or mortality, known only from laboratory experiments (*Pseudomonas sp*. infecting *Solenopsis sp*., Table S2).

### 4.5 Fungi

Ants have a long evolutionary history with fungi (Chapela *et al*., 1994; Hajek & St. Leger, 1994; Mueller, Rehner, & Schultz, 1998). The symbiosis between leaf-cutter ants and their fungi dates back at least 50 million years ago (Chapela *et al*., 1994; Mueller *et al*., 1998; Aanen *et al*., 2002; Schultz & Brady, 2008), and records of ant parasitism by fungi have been preserved in leaf fossils from 48 million years ago (Hughes, Wappler, & Labandeira, 2010). The ecological interactions between ants and fungi are diverse, ranging from mutualism (e.g. between leaf-cutting ants and their cultivated fungi), to facultative parasitism (e.g. by generalist entomopathogens) and parasitoidism (e.g. by the zombie ant fungus *Ophiocordyceps*).

#### 4.5.1 Fungi as pathogens and parasites of ants

The diversity and pathology of fungi infecting insects has been extensively reviewed by (Madelin, 1966; Hajek & St. Leger, 1994; Evans, 2001; Espadaler & Santamaria, 2012; Araújo & Hughes, 2016; Butt *et al*., 2016). It is outside the scope of this present work to review the fungal biology involved; here we provide a brief overview of each family of fungi recorded infecting individual ants and the records thereof. We do not include records of fungi found in or around ant colonies (e.g. fungi isolated from soil or leaf litter near colonies, from colony refuse piles, or from ant fungal farms), nor do we include records where fungi have been isolated from ant external surfaces without indication of pathology. Interested readers are referred to Baird et al. (Baird *et al*., 2007), Reber and Chapuisat (Reber & Chapuisat, 2012), Woolfolk et al. (Woolfolk *et al*., 2016a) and references therein for surveys of fungi generally associated with ant colonies. We present our records of fungi known to infect ants in Table S2.

Fungal epidemics in ant colonies are rarely reported in the literature (reviewed by (Bequaert, 1922; Schmid-Hempel, 1998; Evans, 2001; Espadaler & Santamaria, 2012). As Espadaler and Santamaria note in their review of ecto- and endoparasite fungi of ants: “Extensive, massive mycoses are an extremely rare instance in ants (Hölldobler & Wilson, 1990) and involve individuals, rather than whole colonies”. Evans (Evans, 1974) notes: “Fungal pathogens of ants and other arthropods have been regularly collected from Ghanaian cocoa farms (H.C. Evans, unpublished) but epizootics are of infrequent occurrence and disease is at an enzootic level”. Many fungi have been isolated from in and around ant colonies (Baird *et al*., 2007; Reber & Chapuisat, 2012; Angelone & Bidochka, 2018) and from ant internal tissues (Woolfolk *et al*., 2016a), often for the purposes of identifying potential biocontrol agents, but these have not been accompanied by any observations of colony collapse due to disease. Indeed, in a long-term study monitoring colonies infected by the behaviorally-manipulating fungus *Ophiocordyceps unilateralis* in the field, Loreto et al. (Loreto *et al*., 2014) note that all 17 surveyed colonies continued to function over the 20-month observational period despite chronic infection by a fungal disease.

The absence of fungal disease epizootics in natural ant populations contrasts starkly to the reported impacts of fungal disease on ant colonies in laboratory settings, where generalist fungal pathogens in the genera *Metarhizium* and *Beauveria* are routinely used to investigate potential mechanisms of social immunity (reviewed in (Loreto & Hughes, 2016)). Large infective doses and the use of fungal pathogens which are not naturally co-evolved with ants might explain the discrepancy between fungal-induced mortality in the lab and the paucity thereof in the field (Loreto & Hughes, 2016).

#### 4.5.2 Records of fungi infecting ants [362 records]

We report 362 records of parasitic fungi (9 families, 17 genera, 75 species) infecting 52 ant genera and 239 ant species (Table S2). These fungi come from two phyla (Ascomycota and Entomophthoromycota) and 5 orders (Entomophthorales, Eurotiales, Hypocreales, Laboulbeniales, and Pleosporales).

#### 4.5.3 Ancylistaceae (Entomophthoromycota: Order Entomophthorales) [1 record]

One species of fungi in the family Ancylistaceae has been found parasitizing ants *(Conidiobolus* infecting *Solenopsis invicta*, (Sanchez-Peña & Thorvilson, 1992)). *Conidiobolus* is a pathogen of termites (Kevorkian, 1937; Yendol & Paschke, 1965) and aphids and very occasionally infects vertebrates (Humber, Brown, & Kornegay, 1989). In the case of ants, *Conidiobolus* was recorded from 3 out of approximately 1,000 samples of recently mated red imported fire ant queens (Sanchez-Peña & Thorvilson, 1992).

#### 4.5.4 Entomophthoraceae (Entomophthoromycota: Order Entomophthorales) [13 records]

We report the genera *Tarichium* and *Pandora* as the only Entomophthoraceae fungi known to infect ants, although *Batkoa* has recently been discovered infecting a ponerine ant in South Africa (J. Araujo and D. Hughes, unpublished data).

*Tarichium* has only been reported from ants once (infecting *Tetramorium caespitum* in Russia, (Marikovsky, 1962), but it is an intriguing record as it appears to have caused a “dying out of a large number of this species”. Marikovsky notes that healthy ants began emigrating from infected colonies, perhaps to escape parasite pressure. No other reports of *Tarichium* infecting ants have been observed, nor have other outbreaks killing off whole colonies. The details on the relationship and prevalence of *Tarichium* infecting ants are unfortunately scarce, and follow-up work is needed to confirm the veracity of the fungal identity. Due to the similarities in the biology of *Tarichium* and fungi in the genus *Pandora*, this record likely belongs in the genus *Pandora* (Małagocka, Jensen, & Eilenberg, 2017).

*Pandora* (12 records) is reported parasitizing ants exclusively from the genus *Formica* in Europe (reviewed within (Csata *et al*., 2013)). The genus currently consists of 41 species, of which two have been reported infecting ants (*P. formicae* and *P. myrmecophaga*), though their status as separate species is unclear (Małagocka *et al*., 2017). This fungus causes the characteristic symptoms of summit disease, well-described in (Marikovsky, 1962; Boer, 2008). As noted by Malagocka et al. (Małagocka *et al*., 2017) and observed in other specialized fungal entomopathogens (Hughes *et al*., 2011; Loreto *et al*., 2014), the fungus is unlikely to sporulate inside the nest, and must grow and therefore be transmitted outside of ant colonies.

#### 4.5.5 Trichocomaceae (Ascomycota: Order Eurotiales) [8 records]

Generalist fungi in the family Trichocomaceae belonging to the genera *Aspergillus, Neosartorya, Paecilomyces, Penicillium*, and *Petromyces* have been recorded infecting ants. The majority of these records are for ants in the genera *Solenopsis* and *Paratrechina*, where targeted surveys have actively searched for fungal pathogens as potential use as biocontrol agents against these pestiferous genera (Rodrigues *et al*., 2010; Woolfolk *et al*., 2016a). Other ant genera infected include *Formica* and *Atta*.

For several of these records, the fungus was isolated from ant internal tissues in the absence of ants exhibiting any pathological symptoms (i.e. records within (Woolfolk *et al*., 2016a), so the morbidity and mortality caused by these fungi remains unknown. Other fungi within this family have been demonstrated to infect and kill their ant hosts (i.e. species listed within (Fernández-Marín *et al*., 2006; Rodrigues *et al*., 2010)); however, these were laboratory infections and there are no records of colonies naturally infected by these fungi in the field. Evans notes that *Aspergillus* and *Penicillium* have been associated with moribund ants, but considers them to be “opportunistic pathogens on damaged or stressed ants” (Evans, 2001).

#### 4.5.6 Pleosporaceae (Ascomycota: Order Pleosporales) [1 record]

*Alternaria tenuis* infecting *Formica rufa* (Marikovsky, 1962) is an *ex-nido* transmitting fungal parasite, requiring the death of its ant host outside of the nest in order to be able to develop and properly transmit. Infected ants undergo behavioral changes collectively referred to as ‘summit disease’ (Marikovsky, 1962; Boer, 2008). Marikovsky reports that disease due to *Alternaria* occurred primarily in the late summer or early fall, and that fungal development on the outside of the host, and thus onward transmission, was prevented if the death of the host occurred on particularly dry or sunny days. Evans (2001) notes that the identification of the fungus as *Alternaria* might be doubtful, because *“A. tenuis* is a common saprophyte and has no previous or present history as an entomopathogen. Indeed, the host and the description of the behaviour and habitat suggest that the true pathogen is in fact a member of the Entomophthoraceae (Bałazy & Sokołowski, 1977) and that *A. tenuis* is an opportunistic invader, possibly outcompeting the primary pathogen.” (Evans, 2001).

#### 4.5.7 Laboulbeniaceae (Ascomycota: Order Laboulbeniales) [39 records]

We report 39 records of fungi from the family Laboulbeniaceae associated with ants. These records represent only 2 genera, *Laboulbenia* and *Rickia*, primarily infecting the ant genera *Formica, Lasius*, and *Camponotus* in Europe and the United States.

The biology and host records of *Laboulbenia* and *Rickia* are reviewed in Espadaler and Santamaria (2012); both fungi are obligate ectoparasites, growing on the cuticle of ants without appearing to harm the host (Espadaler & Santamaria, 2012). Transmission is presumably accomplished by direct contact with fungal material between nest mates. Evans (2001) notes that the Laboulbeniaceae have never been implicated in host mortality and are “commensal parasites” of ants. In our database, we note them as parasites, but clearly more studies are needed to determine their impact on individual ant health.

#### 4.5.8 Clavicipitaceae (Ascomycota: Order Hypocreales) [7 records]

*Metarhizium* is the only genus of fungi within the Clavicipitaceae reported to infect ants. This generalist fungus requires the death of its insect host. Following death, hyphae emerge from the insect body and sporulate, and transmission occurs via direct contact with spores. This genus is incredibly cosmopolitan, infecting many species of insects (Vega & Blackwell, 2005), and it appears to be an important part of the soil microbial community. It was recently found to be common in soil around ant nests in Ontario, Canada (Angelone & Bidochka, 2018).

While this fungus is used extensively in the laboratory to investigate mechanisms of ant social immunity (reviewed by (Loreto & Hughes, 2016) and despite its wide distribution in soil microbial communities (Leger, 2008), it has been rarely reported infecting ants in natural populations (reviewed in (Evans, 2001; Loreto & Hughes, 2016)). The first recorded instance of *Metarhizium* infecting an ant is from Petch (Petch, 1931), who noted it infecting a “black ant”. It has since been reported from *Atta, Formica, Leptogenys, Paratrechina*, and *Solenopsis*. There are no reports of ant colony epizootics due to *Metarhizium* in natural populations.

#### 4.5.9 Cordycipitaceae (Ascomycota: Order Hypocreales)

[31 records]

The Cordycipitaceae were previously grouped within the Clavicipitaceae; revision by Sung et al. (2007) erected the Cordycipitaceae as its own family (Sung *et al*., 2007). Two genera within the Cordycipitaceae are known as parasites of ant hosts: *Akanthomyces* and *Beauveria*.

*Akanthomyces gracilis* is reported from *Camponotus, Crematogaster Dorylus, Macromischoides, Oecophylla, Paltothyreus, Platythyrea*, and *Polyrhachis* (reviewed in (Evans, 2001)). Evans (2001) notes that when dead insects infected by this fungus are hidden in the soil or leaf litter, they bear long synnemata “which ramify above or within the substrate”. However, it is unclear whether *Akanthomyces* is the causative agent of host death, or merely an opportunistic hyperparasite (J. Araújo, personal communication). Additional studies are needed to ascertain its relationship to the ant host.

*Beauveria* has been found to naturally infect over 700 insect species and has been used prolifically for the purposes of biocontrol (reviewed in (Inglis *et al*., 2001; Meyling & Eilenberg, 2007)). *Beauveria* has recently been found to be paraphyletic (Rehner & Buckley, 2005), so though we report a single species, *Beauveria bassiana*, infecting ants, we can expect additional records as the diversity within *Beauveria* is unraveled. *B. bassiana* has been found infecting ants in the genera *Atta, Camponotus, Cephalotes, Crematogaster, Ectatomma, Formica, Lasius, Myrmecia, Oecophylla, Paraponera*, and *Solenopsis*. Like *Metarhizium, Beauveria* has been used as a laboratory workhorse to understand potential mechanisms of social immunity in ant colonies (Loreto & Hughes, 2016). Epizootics due to *Beauveria* in natural ant populations are unreported.

#### 4.5.10 Ophiocordycipitaceae (Ascomycota: Order Hypocreales) [197 records]

The majority of records of fungi infecting ants come from the family Ophiocordycipitaceae. Prior to its re-erection in 2007, members of the Ophiocordycipitaceae had been placed within the polyphyletic family Clavicipitaceae, genus *Cordyceps*. The Ophicordycipitaeceae contains the striking ‘zombie ant’ fungus *Ophiocordyceps*, known for the sophisticated and species-specific behavioral manipulation of its host (Hughes *et al*., 2011; Andersen *et al*., 2012; de Bekker *et al*., 2014, 2015; Fredericksen *et al*., 2017; Araújo *et al*., 2018). Prior to the adoption of the ‘One Fungus, One Name’ policy (Taylor, 2011), the genera *Hirsutella, Hymenostilbe, Stillbella*, and *Stilbum* were also included in this family, but these are asexual stages (= anamorphs) of *Ophiocordyceps* and thus are no longer used (Spatafora *et al*., 2015). Other genera infecting ants in this family are *Paraisaria* and *Polycephalomyces*.

*Ophiocordyceps* is an old genus that had been moved to be within *Cordyceps* but was re-erected following the major revision of the Clavicipitaceae (Sung *et al*., 2007; Quandt *et al*., 2014). There has been much focus on this genus in recent years, with many new species being described from former species complexes (Evans, Elliot, & Hughes, 2011; Araújo *et al*., 2015, 2018). These species complexes, including *O. unilateralis s.l., O. australis s.l., O. kniphofioides s.l., O. myrmecophila s.l*., and *O. sphecocephala s.l*., undoubtedly contain many new species awaiting description. With the exception of *O. sphecocephala s.l*., all are parasitic on ants.

#### 4.5.11 Incertae Sedis [65 records]

Four genera of fungi infecting ants are currently without family-level placement: *Aerigitella, Hormiscium, Myrmecomyes*, and *Mymicinosporidium*.

*Aerigitella* (23 records) is an ectoparasitic fungus known primarily from *Formica* and *Lasius* in Europe and the United States (Espadaler & Santamaria, 2012). As reviewed by Espadaler and Santamaria (2012), the relationship of this fungus with its ant host, its life history, and mode of transmission are unknown. Some suggestions that it might cause reduced life span or activity have been noted (Chérix, 1982; Wisniewski & Buschinger, 1982).

*Hormiscium* (3 records) is an ectoparasitic fungus described from *Pseudomyrmex* and *Myrmica* in South America and Europe, respectively (reviewed in (Espadaler & Santamaria, 2012)). Little information is known about its biology and relationship to its ant host.

*Myrmecomyces* (1 record) is reported infecting *Solenopsis* ((Jouvenaz & Kimbrough, 1991), within (Evans, 2001). Its effects on host morbidity and mortality are unknown.

*Myrmicinosporidium durum* (39 records) is an endoparasitic fungus of uncertain phylogenetic position that has been found infecting many different ant genera and species (Sanchez-Peña, Buschinger, & Humber, 1993; Pereira, 2004; Csősz *et al*., 2012; Espadaler & Santamaria, 2012; Gonçalves, Patanita, & Espadaler, 2012). Though first described in the 1920's (Hölldobler, 1927, 1933), the life-history of the fungus remains unknown, and it is unclear what the effect of infection does to the ant host in terms of either morbidity or mortality (Sanchez-Peña & Thorvilson, 1992; Espadaler & Santamaria, 2012). Sanchez-Pena et al. (1993) note that the fungus “does not seem to impair significantly the mobility or behavior of the ant hosts, despite extensive vegetative development and sporulation prior to host death”. There has been discussion over whether *Myrmicinosporidium* might be a single generalist species (Gonçalves *et al*., 2012), or whether it could constitute a fungal complex with many new species awaiting description (Pereira, 2004; Espadaler & Santamaria, 2012). The enigmatic life history and host relationship of this fungus, coupled with its wide host range and geographic distribution make this fungus ripe for future work.

### 4.6 Microsporidia and Apicomplexa

Microsporidia are a group of obligate, unicellular, spore-forming parasites (Keeling & Fast, 2002). Historically, microsporidians have been grouped within the protozoa, but they have been found to be closely related to fungi (Hirt *et al*., 1999), though their larger taxonomic placement (i.e. within the fungi or as a sister group to fungi) remains uncertain. One family within the Microsporidia, Burenellidae, has been recorded infecting ants from the genus *Solenopsis*, though some parasites in the group are awaiting family-level placement. We report 13 records of microsporidian and apicomplexan parasites infecting ants.

#### 4.6.1 Microsporidia: Burenellidae [5 records]

Two genera within the Burenellidae, *Burenella* and *Vairimorpha*, have been found infecting *Solenopsis* in South America. *Burenella dimorpha* was first described by Jouvenaz and Hazard (1978) infecting the fire ant *Solenopsis geminata* (Jouvenaz & Hazard, 1978). As noted by those authors, host pathology is only observed in pupae, and infected pupa have been observed being cannibalized in the laboratory (Jouvenaz, Lofgren, & Allen, 1981). Transmission experiments show that the parasite is transmitted to 4th-instar larvae *per os* by workers tending brood (Jouvenaz & Hazard, 1978; Jouvenaz *et al*., 1981). Adults don't seem to become infected because they are able to filter out infective spores in their infrabuccal pocket (Jouvenaz *et al*., 1981).

*Vairimorpha* is a closely related genus also found infecting *Solenopsis*. Unlike *Burenella*, Jouvenaz and Ellis (1986) found that all ant stages of S. *invicta* (i.e. brood and adults) could be infected, but do not report any physical or behavioral symptoms associated with infection (Jouvenaz & Ellis, 1986). Briano et al. (2002) surveyed many ant genera for the presence of microsporidians, but only report finding *Vairimorpha-i*nfected Solenopsis (S. invicta, S. *macdonaghi*, and S. *richerti)* at low prevalences (Briano *et al*., 2002).

#### 4.6.2 Microsporidia: *Incertae Sedis* [4 records]

*Kneallhazia solenopsae* (formerly *Thelohania solenopsae*, in the family Thelohaniidae) was first described from *Solenopsis invicta* in Brazil (Allen & Buren, 1974). Surveys have found it to be the most common microsporidian parasite infecting fire ants in South America (Allen & Silveira-Guido, 1974; Jouvenaz *et al*., 1981; Briano, Patterson, & Cordo, 1995; Briano *et al*., 2002), and it has also been found parasitizing some S. *invicta* colonies in the southeastern United States (Williams, Oi, & Knue, 1999). Sokolova and Fuxa (2008) found that spores can be transmitted both transovarially and *per os* through trophallaxis, infecting all stages from brood to adults (Sokolova & Fuxa, 2008). In both brood and adults, infection related pathology is not apparent (Sokolova & Fuxa, 2008), though Allen and Buren (1974) noted that infected colonies had a “noticeable loss of vigor and pursuit when disturbed” (Allen & Buren, 1974).

#### 4.6.3 Apicomplexa: Lipotrophidae [4 records]

One genus from the Apicomplexa, *Mattesia*, is recorded infecting species of *Solenopsis* in Brazil and the United States (Jouvenaz *et al*., 1980; Pereira *et al*., 2002). Infections by *Mattesia* are apparent in brood, which eventually become almost solid black in coloration and do not complete their development (Jouvenaz & Anthony, 1979). Infected adult ants have not been observed (Jouvenaz & Anthony, 1979; Kleespies *et al*., 1997; Pereira *et al*., 2002), perhaps due to the filtering of infective spores through their infrabuccal pocket.

### 4.7 Trematoda

Trematodes (‘flukes', phylum Platyhelminthes) are a class of flatworms with a wide diversity of free-living and parasitic lifestyles (Esch, Barger, & Fellis, 2002). Parasitic trematodes use multi-host life cycles involving two to three hosts (Poulin, 1998; Bush, 2001; Esch *et al*., 2002), in which mollusks serve as the first intermediate hosts and vertebrates serve as final hosts. Trematodes infecting ants are few but spectacular, and trematodes from two families, the Dicrocoelidae and the Heterophyidae, are recorded infecting ants. In these instances, the ants serve as second intermediate hosts in the multi-host life cycles of these trematodes. We report 21 records of trematodes from two families infecting five genera of ants (Table S2).

#### 4.7.1 Dicrocoeliidae [20 records]

Two genera, *Brachylecithum* and *Dicrocoelium* are reported infecting ants. *Brachylecithum* is reported from *Camponotus* in the United States (Carney, 1967, 1969), and *Dicrocoelium* is reported infecting *Cataglyphis, Camponotus, Formica*, and *Lasius* in Europe and Asia (reviewed in (Manga-González *et al*., 2001)). The life cycle of both parasites is complex, involving snails as first intermediate hosts and vertebrates as final hosts, and it took over 100 years before the life cycle was elucidated by Krull and Mapes (1952) (Krull & Mapes, 1952). Infected land snails produce slime balls containing trematode cercariae, which ants readily eat. The cercariae then penetrate the ant's crop (social stomach) and enter the hemocoel, where they migrate through the ant's body to the gaster and brain, forming metacercarial cysts (Carney, 1969). Carney (1969) found that it could take as long as 120 days for the metacercarial cysts to develop and become infective. Once the trematodes are ready to find their next host, infected ants display several behavioral changes that serve to increase the probability of trophic transmission to the next host. These behavioral changes include sluggishness, lack of sensitivity to light, and more time spent outside the nest (see description in (Carney, 1969)).

#### 4.7.2 Heterophyidae [1 record]

*Eurytrema pancreaticum* is reported infecting *Technomyrmex detorquens* (Svadzhyan & Frokolva, 1966). The life cycle of this parasite is thought to be similar to that of dicrocoelid trematodes infecting ants, though due to the limited number of records, any behavioral changes in infected ants associated with this parasite are unknown.

### 4.8 Cestoda

Cestodes (‘tapeworms', phylum Platyhelminthes) are a class of parasitic flatworms that typically live in the digestive tracts of vertebrate final hosts. Ants are intermediate hosts in the life cycles of several cestode genera from three families: Davaineidae, Dilepidae, and Mesocestoididae. Cestode-infected ants have been recorded primarily due the impact that these tapeworms have on their final hosts (i.e. poultry and other domesticated animals, occasionally including humans). We report a total of 66 records of cestodes infecting ants, encompassing three families of cestodes infecting 44 ant species (Table S2).

Life cycles of cestode-infected ants are fairly straightforward: adult ants find cestode proglottids while foraging and bring these proglottids back into the colony. Proglottids or oncospheres are fed to the brood, which then develop into cysticercoids or procercoids. When infected brood have matured into adults, they are behaviorally more sluggish than their non-infected counterparts, and thus more likely to be consumed by predators who act as second intermediate hosts or final hosts. Following reproduction, the final host deposits cestode-egg-laden feces, which are consumed by ants to begin the cycle anew.

#### 4.8.1 Davaineidae [62 records]

Davaineid cestodes from the genera *Cotugnia* (6 records) and *Raillietina* (56 records) have been recorded infecting ants. *Cotugnia digonopora* has been found infecting *Monomorium* and *Pheidole. Raillietina* (17 species) has been found to infect 14 ant genera in Africa, Asia, Australia, Europe, and North America.

#### 4.8.2 Dilepididae [2 records]

*Anomotaenia brevis* and *Choanotaenia crateriformis* have been recorded from *Lepthorax* in France and Spain.

#### 4.8.3 Mesocestoididae [2 records]

*Mesocestoides* has been occasionally reported infecting humans and domesticated animals, but the life cycle of this tapeworm has remained enigmatic for over a century. It has been thought that oribatid mites and other arthropods could serve as first intermediate hosts, but surveys for the tapeworm in these arthropods have been elusive. Very recently, ants (*Lasius niger* and *Tapinoma sessile*) have been identified as first intermediate hosts in the life cycle (Padgett & Boyce, 2007).

### 4.9 Nematoda

Nematodes (‘roundworms') are a diverse phylum in which more than half of its members use a parasitic lifestyle (Blaxter, 2003; Blaxter & Koutsovoulos, 2015). Parasitism has evolved at least 15 times independently within this phylum over the course of its evolutionary history (Blaxter & Koutsovoulos, 2015). Fossil evidence indicates that relationships between ants and nematodes date back at least 40 million years (Poinar Jr, 2002, 2012; Poinar Jr *et al*., 2006). We report 67 records of parasitic nematodes comprising 10 families and 27 genera infecting 20 ant genera and 43 ant species (Table S2).

The biology, extant knowledge, and host records of nematode parasites of ants have been well reviewed by Poinar (2012) (Poinar, 2012). Here we report known records of nematodes infecting ants and summarize some interesting aspects of their biology. A large number of nematodes in the family Rhabditidae are phoretically associated with ants (see SI), and the majority of nematodes infecting ants belong to the family Mermithidae.

#### 4.9.1 Allantonematidae [2 records]

The Allantonematidae reported infecting ants comprise two genera: *Formicitylenchus* (extant) and *Paleoallantonema* (fossil). *Formicitylenchus oregonensis* is recorded from *Camponotus vicinus* in the United States (Poinar, 2003). Poinar (2012) notes that while many life cycle details remain unknown for this parasite, it is likely that free-living worms penetrate the cuticle of ant larvae and develop as the ant develops (Poinar Jr, 2012). Poinar (2012) suggests that the nematode is transmitted by infected queen ants but impacts on host behavior and morbidity or mortality are unknown. *Paleoallantonema cephalotae* is described infecting *Cephalotes serratus* from a fossil in Dominican amber (Poinar, 2011), suggesting that the relationship between allantonematids and ants may be more widespread than what our current host records suggest (Poinar Jr, 2012).

#### 4.9.2 Diplogastridae [2 records]

The Diplogastridae, along with the Rhabditidae and Panagrolaimidae, are families of nematodes in which the juvenile stages live phoretically with ants, either inside the pharyngeal glands or outside on the ant's cuticle. The damage to the host caused by these associations is considered minor, if damage even occurs (Markin & McCoy, 1968; Poinar, 2012). Within the Diplogastridae, *Formicodiplogaster myrmenema* (fossil) and *Pristonchus sp*. are recorded from *Azteca alpha* and *Atta cephalotes*, respectively.

#### 4.9.3 Panagrolaimidae [2 records]

One genus in the Panagrolaimidae, *Panagrolaimus*, has been found infecting *Acromyrmex crassispinus* in Paraguay and *Atta cephalotes* in French Guiana (Sudhaus, 2016). Like the Rhabditidae and Diplogastridae, juvenile nematodes from this family are phoretic associates with ants causing minimal damage to their host (Poinar, 2012).

#### 4.9.4 Rhabditidae [29 records]

The Rhabditidae, along with the Displogastridae and Panagrolaimidae, are phoretic associates of ants that cause minimal damage to their host (Poinar, 2012). Many Rhabditidae live within the ant nest environs (see (Sudhaus, 2016)); we only include those records that were isolated from ant bodies (either internal or external, head, body). Nine genera from the Rhabditidae, *Diplogasteroides, Diploscapter, Halicephalobus, Koernia, Mesorhabditis, Oscheuis, Pristionchus, Rhabditis*, and *Sclerorhabditis* are phoretic parasites of 13 genera of ants (Poinar, 2012).

#### 4.9.5 Heterorhabditae [2 records]

The Heterorhabditidae, along with the Steinernematidae, are entomopathogenic nematodes with interesting bacterial associations (Poinar, 2012). The nematodes infect ants through the transmission of food and, after gaining entrance to the hemocoel, release symbiotic bacteria that kill the ant host. Records of nematodes from this family infecting ants are known only from the laboratory; no natural infections have yet been observed. *Heterorhabditis bacteriophora* has been found in association with *Solenopsis invicta* and S. *richteri* ((Quattlebaum, 1980), within (Poinar, 2012)).

#### 4.9.6 Steinernematidae [8 records]

The Steinernematidae, along with the Heterorhabditidae, use symbiotic bacteria to kill their ant host after entering the hemocoel following trophallaxis (Poinar, 2012). Though only known from laboratory infections, *Steinerema carpocapsae* has been recorded successfully infecting ants from the genera *Acromyrmex, Camponotus, Myrmica, Pogonomyrmex*, and *Solenopsis* (see review within (Poinar, 2012)).

#### 4.9.7 Physalopteridae [1 record]

*Skrjabinoptera phrynosoma* is the sole species of this family recorded infecting ants. The final hosts of this nematode are lizards, who excrete nematodes containing infective eggs in their feces. Foraging ants feed these nematodes and their eggs to brood, wherein the nematode eggs develop into larval and then juvenile stages as the ant develops through eclosion. When lizards eat the infected adult ants, the life cycle is completed. Poinar (2012) notes that infected workers containing more than ten nematodes were still active, but had enlarged, lighter-colored gasters (Poinar, 2012).

#### 4.9.8 Seuratidae [1 record]

Only one record of a nematode from the family Seuratidae, normally known to infect vertebrates, has been found infecting ants. *Rabbium paradoxus* was found infecting *Camponotus castaneus* in the United States (Poinar, Chabaud, & Bain, 1989). It is thought that ants became infected after ingesting nematode eggs contained within lizard feces. Poinar (2012) notes that parasitized workers had swollen gasters and were found foraging during the day, which would make them easier targets for predation and thus trophic transmission of the nematode (Poinar, 2012).

#### 4.9.9 Tetradonematidae [3 records]

We report three records of Tetradonematids infecting ants from two genera: *Tetradonema* and *Myrmeconema. Tetradonema solenopsis* has been found parasitizing *Solenopsis invicta* in Brazil ((Nickle & Ayre, 1966), within (Poinar, 2012)); most life cycle details and the impact of infection due to these nematodes remain unknown (Poinar, 2012).

*Myrmeconema* is a genus consisting of two species *(M. antiqua-* fossil, *M. neotropicum-* extant) that make use of trophic transmission in order to complete their life cycle (Yanoviak *et al*., 2008; Poinar, 2012). Ant pupae are fed female nematodes; once the ant becomes an adult, nematode egg masses are released into the ant's hemocoel and change the color of the ant's gaster from black to red. Infected ants hold their gasters high in the air, which combined with the red coloration of their gasters, makes them appear to be fruit. Birds eat the ants and the nematode finishes its life cycle inside its final host. Though *M. antiqua* is a fossil species and thus its life cycle details cannot be definitively described, it is thought that its life history is very similar to that of *M. neotropicum* (Poinar, 2012). *M. neotropicum* is a striking example of combined behavioral and morphological manipulation of a host by a parasite.

#### 4.9.10 Mermithidae [17 records]

Mermithid nematodes are well-known parasites of ants due to the stunning examples of behavioral manipulation that they cause (reviewed in (Moore, 2002)). While many mermithid species reported from ants are undescribed (see (Poinar Jr *et al*., 2006)), we report eight genera (1 fossil, 7 extant) infecting ants: *Agamomermis*, *Allomermis*, *Camponotimermis*, *Comanimermis*, *Heydenius* (fossil), *Mermis, Meximermis*, and *Pheromermis*.

The general life cycle of a mermithid nematode is reviewed in Poinar (2012) (Poinar Jr, 2002). Foraging ants collect protein sources that serve as paratenic hosts for the nematode (e.g. the aquatic larvae of dipterans). These paratenic hosts are fed to ant brood wherein the nematode initiates development. Once the nematode has finished developing inside the now adult ant, it induces its host to seek out water. Once there, the adult nematode escapes its dying ant host to complete the remainder of its life cycle in aquatic, larval-stage insects.

### 4.10 Hymenoptera

Hymenoptera are a large, diverse order of insects containing the ants, bees, wasps, and sawflies. Recently, it has been suggested that hymenopterans may be even more diverse than beetles, thought to be the most species-rich animal order, primarily due to the large number of parasitoid species (Forbes *et al*., 2018). Hymenoptera have a wide range of lifestyles, from solitary to eusocial, and free-living to parasitoid. The parasitoid lifestyle in the Hymenoptera had a single origin early in the Euhymenopteran lineage (Whitfield, 1998, 2003), with ectoparasitoidism as the ancestral mode. Many independent transitions to endoparasitism have occurred within parasitoid lineages (Whitfield, 1998; Murray, Carmichael, & Heraty, 2013). Interestingly, the evolution of complex brain structures known as mushroom bodies was concurrent with the acquisition of parasitoidism and not with the rise of sociality as previously thought (Farris & Schulmeister, 2011). Interested readers should refer to Whitfield (1998, 2003) and Eggleton and Belshaw (1992) for reviews of hymenopterans as parasitoids (Eggleton & Belshaw, 1992; Whitfield, 1998, 2003).

Hymenopteran parasitoids of ants have been well-reviewed by Lachaud and Perez-Lachaud (2012) and by Schmid-Hempel (1998) (Schmid-Hempel, 1998; Lachaud & Pérez-Lachaud, 2012). We summarize interesting aspects of parasite biology for each hymenopteran family below and report 355 records of Hymenoptera from 11 families infecting ants (Table S2). The majority of these records come from the Eucharitidae, which are ecto- and endoparasitoids of ant brood, followed by the Braconidae, Diapriidae, and Encyrtidae.

#### 4.10.1 Mutillidae [5 records]

The Mutillidae are a large family of wasps commonly known as ‘velvet ants'. Mutilid larvae are idiobiont ectoparasitoids of enclosed host stages (e.g. pupae) (Brothers, Tschuch, & Burger, 2000). In many cases, direct parasitism of ants by mutillids has not been observed, but the association is inferred (Brothers, 1994). Many mutillids are parasitoids of commensals living inside ant nests (e.g. beetles) and are not parasitoids of the ants themselves. In other instances, mutillids are Mullerian mimics of ants (Brothers *et al*., 2000). In a recent revision of the mutillid subfamily Rhopalomutillinae, Brothers (2015) noted that the biology of the wasps indicated that they might parasitize ants (Brothers, 2015). More work is needed to determine the extent of host-parasite relationships between the Mutillidae and ants.

While Mutillidae commonly parasitize solitary bees and wasps (Brothers, 1989; Brothers *et al*., 2000), only five species within the genus *Ponerotilla* have been reported (presumably) parasitizing immature workers of the ponerine ant *Brachyponera lutea* in Australia ((Brothers, 1994), within (Schmid-Hempel, 1998). The general life cycle of mutillids parasitizing insects seems to be that following mating, female mutillids independently enter insect nests and deposit eggs near larvae and pupae. Developing wasps then feed and kill their larval/pupal hosts as ectoparasitoids.

#### 4.10.2 Chalcididae (Superfamily Chalcidoidea) [5 records]

Chalcidid wasps are endoparasitoids (occasionally ectoparasitoids) of Diptera and Lepidoptera, though other orders of insects, including Hymenoptera, may be infected (Noyes, 2017). We report five species of chalcidid wasps from the genus *Smicromorpha* infecting ants, though some of these reports have not been conclusively confirmed (see discussion in (Darling, 2009)). All are parasitoids of the arboreal weaver ant, *Oecophylla smaragdina*, in the Indo-Australia region (Darling, 2009; Lachaud & Pérez-Lachaud, 2012; Noyes, 2017). Weaver ants use their silk-spinning larvae to create their leaf nests in trees; *Smicromorpha* deposits eggs on the larvae while they are exposed during the nest building process ((Girault, 1913), within (Darling, 2009)). It appears that the association between *Smicromorpha* and *Oecophylla* is unique, and given the overlapping distributions of both these genera in mainland Southeast Asia, reports from that area should be expected as more surveys are conducted (Darling, 2009).

#### 4.10.3 Encyrtidae (Superfamily Chalcidoidea) [1 record]

The Encyrtidae are a family of wasps that are endoparasitoids or egg predators primarily of scale insects, but also other orders of insects (Noyes, 2017). Many encyrtid wasps are reported as associates of ants, but few of these are actually parasitic. As reviewed in Perez-Lachaud, Noyes, and Lachaud (2012):

> “Numerous cases of associations of encyrtid wasps with ants have already been reported. In the majority of these cases, however, wasps are associated only indirectly with ants (interference associations) through primary parasitism of the trophobionts (Coccoidea), which are exploited and protected by ants. Suspected direct parasitism cases are unusual, and no direct attack of encyrtids on ants has ever been demonstrated.”.

Only recently has an encyrtid been found as a true primary parasite of ants. A single species, *Blanchardiscus pollux*, is reported infecting the ponerine ant *Neoponera goeldii* in French Guiana (host species is listed as *Pachycondyla goeldii*, (Pérez-Lachaud, Noyes, & Lachaud, 2012)). More surveying might reveal additional encyrtid wasps that are primary parasites of ants.

#### 4.10.4 Eucharitidae (Superfamily Chalcidoidea) [199 records]

Wasps within the family Eucharitidae are parasitoids of ant brood including larvae, pre-pupae, and pupae (Clausen, 1940; Heraty, 2002; Heraty *et al*., 2004; Heraty, Heraty, & Torréns, 2009; Noyes, 2017). Eucharitid wasps colonized the ants approximately 72 million years ago (Murray *et al*., 2013) and are known worldwide (Heraty, 2002, 2014). Murray et al. (2013) provide a wonderful study on the phylogeny of Eucharitidae as it relates to that of their ant hosts, “Eucharitidae exhibit a general trend of ant subfamily colonization (host-switching) occurring infrequently at an early time period, followed by high host conservatism (phylogenetic affinity) at the ant-subfamily level in extant lineages” (Murray *et al*., 2013).

We report 199 records encompassing 109 eucharitid species in 29 genera as parasitoids of 30 genera of ants. Many additional eucharitid species have been described, but full life history details including their associated host ant species remain unknown (see (Torréns, 2013; Heraty, 2014)). It can be expected that as these life history details are filled in, numerous host records will be added to the database.

The general life cycle of eucharitid wasps can be described as follows: adult female eucharitid wasps locate the microhabitat of potential hosts and lay eggs on nearby vegetation, which can include leaves and fruit (Heraty, 2014). These eggs develop into first-instar planidial larvae (described by (Bouček, 1956; Heraty & Darling, 1984)), which must then make contact with the ant host (Heraty, 2002; Lachaud & Pérez-Lachaud, 2012). It is currently unknown whether the larvae actively seek out potential hosts, or are picked up through passive contact with foraging ants (Noyes, 2017). Inside the nest, the planidial larvae need to gain access to the brood, which is accomplished through phoretic transfer from workers tending the brood.

Once they have accessed ant larvae, eucharitid wasps can either be internal or external parasitoids of their ant host (Noyes, 2017), both of which ultimately result in host death. It appears that external parasitism (ectoparasitism) is the most likely ancestral state, with internal parasitism (endoparasitism) having evolved independently in at least three lineages of the Eucharidtidae (Heraty & Murray, 2013). As either ecto- or endoparasites, eucharitid larvae generally wait until the host has reached the pre-pupal stage before continuing their larval development and can pupate inside the host cocoon or naked inside the nest (Noyes, 2017). Upon completion of their development, adult eucharitid wasps must successfully exit the ant nest in order to find mates. It seems that some eucharitids might exploit the hygienic behavior of their hosts, using imperfectly mimicked host cuticular carbon profiles in order to be identified as intruders and personally escorted out (Pérez-Lachaud *et al*., 2015). How far the adult wasps disperse is not well known, but Ayre (1962) found that mating between *Pseudometagea schwarzii* wasps occurred almost immediately following emergence, and they laid their eggs within the foraging area of their host ant nest (Ayre, 1962).

Despite their worldwide distribution and parasitoidism of many ant genera, eucharitid wasps do not seem to impose a major cost on ant colonies. Ayre (1962) writes, “Though parasitism is very high within the centers of infestation, the influence of *P. schwarzii* on the population size of *L. neoniger* is doubtful, even within these centers.” (Ayre, 1962). This observation has been made by others; Lachaud and Perez-Lachaud note, “Nevertheless, considering the extremely high nest densities recorded in the study zone (Schatz & Lachaud, 2008), the *E. ruidum* population did not appear to be seriously affected by such a high level of parasitism.” (Lachaud & Pérez-Lachaud, 2012).

#### 4.10.5 Eulophidae (Superfamily Chalcidoidea) [13 records]

The Eulophidae are a speciose and globally distributed family of wasps that are primarily parasitoids of concealed larvae (Noyes, 2017). Eulophid wasps can be ecto- or endoparasitoids, soiltary or gregarious, idiobionts or koinobionts of a diverse array of insects (Noyes, 2017).

Many eulophid wasps have been reported as nest associates of ants, attacking commensals that live inside ant nests. We report 13 records wherein the eulophid wasp is a primary parasitoid of ant brood. These ant-parasitic Eulophidae comprise the genera *Melittobia, Horismenus, Myrmokata*, and *Pediobius* infecting the ant genera *Acantholepis, Camponotus, Crematogaster*, and *Formica*.

#### 4.10.6 Eupelmidae (Superfamily Chalcidoidea) [2 records]

The Eupelmidae are a family of wasps that are parasitic on the immature stages of other insects (Noyes, 2017). Only two species of Eupelmidae are brood parasitoids of ants, both within the genus *Anastatus. Anastatus myrmecobius* parasitizes *Temnothorax purpurata*, and *Anastatus reduvii* parasitizes *Pseudomyrmex elongata* (Noyes, 2017).

#### 4.10.7 Eurytomidae (Superfamily Chalcidoidea) [10 records]

The Eurytomidae are a family of wasps with a diverse array of plant-associated life histories, ranging from phytophages to parasitoids of phytophagous insects (Noyes, 2017). We report ten records of eurytomid wasps infecting ants, comprising three species within the genus *Aximopsis*. *Aximopsis affinis* and *Aximopsis aztecicida* parasitize species of *Azteca*, while *Aximopsis sp*. parasitizes *Azteca* as well as a single species of *Camponotus*.

#### 4.10.8 Pteromalidae (Superfamily Chalcidoidea) [2 records]

Only two records of Pteromalidae infecting ants have thus far been reported. *Pheidoloxenus wheeleri* infecting *Pheidole instabilis* and *P. ceres* in the United States and Mexico are the only records to date ((Mann, 1914), within (Schmid-Hempel, 1998)). Mann (1914) writes that W.M. Wheeler, who first discovered the pteromalid infecting ants, considers the pteromalids to be endoparasites of adults or brood (Mann, 1914). Pteromalids have active, host-seeking 1^st^ larval instars, similar to the strepsipterans discussed below, which may be how they access hosts within the colony. The rest of the parasite's biology is unknown.

#### 4.10.9 Braconidae (Superfamily Ichneumonidae) [47 records]

The Braconidae are a very large family of parasitoid wasps, the second most speciose family within the Hymenoptera (Jones *et al*., 2009). Despite this, only four genera are known to infect ants. We report 47 records of braconid wasps in the genera *Elasmosoma, Elasmosomites* (fossil), *Kollasmosoma*, and *Neoneurus* as known parasites of ants. Braconidae infect the Formicinae genera *Formica, Camponotus, Lasius, Polyergus, Cataglyphis*, and the Myrmicine genus *Messor*.

Braconids oviposit directly into adult ants while they are foraging outside of the nest (Forel, 1874; Olivier, 1893; Pierre, 1893; Wasmann, 1897; Donisthorpe, 1927; Kariya, 1932; Durán & van Achterberg, 2011), which is unusual amongst hymenopteran parasitoids (Shaw, 1988). Their larvae develop as endoparasitoids that ultimately kill the ant host (Kistner, 1982; Shaw & Huddleston, 1991; Shaw, 1993; Poinar, 2004). Fossil evidence of a braconid wasp infecting an ant indicates that parasitism by braconids could date back at least 40 million years (Poinar Jr & Miller, 2002).

#### 4.10.10 Ichneumonidae (Superfamily Ichneumonidoidea) [33 records]

The Ichneumonidae are a very large family of parasitic wasps that commonly parasitize the brood of other insects (Yu, 1997). We report three genera of ichneumonid wasps, *Eurypterna, Ghilaromma*, and *Hybrizon* parasitizing ants in the genera *Formica, Lasius, Myrmica*, and *Tapinoma*.

While the life history details and host records for many ichneumonid wasps remain unclear, it appears that Ichneumonidae oviposit directly into ant brood as they are being carried between nests by workers (Komatsu & Konishi, 2010; Durán & van Achterberg, 2011) or in disturbed nests (Marsh, 1989). Following oviposition, the wasps develop as endoparasitoids, ultimately killing their ant hosts (Van Achterberg, 1999).

Komatsu and Kinishi (2010) report an undescribed ichneumonid wasp that took much longer to oviposit in its slower-moving *Myrmica kotokui* host than it took other observed ichneumonids to oviposit in their hosts (Komatsu & Konishi, 2010). They suspect that there might be a diversification in the behavioral ecology of the ichneumonid wasps that reflects how quick-moving their host species are.

#### 4.10.11 Diapriidae (Superorder Diaproidea) [38 records]

The Diapriidae are a family of small parasitic wasps with a worldwide distribution (Masner, 1993). While most Diapriidae attack the larval stages of flies, some are endoparasitoids of ant brood or nest associates living within ant colonies (reviewed in (Loiacono, 1987; Loiácono & Margaria, 2000; Masner & Garcia R, 2002; Lachaud & Pérez-Lachaud, 2012; Loiácono, Margaria, & Aquino, 2013). The life history details for many species are unknown and how they interact behaviorally with their ant hosts is also largely unknown for the majority of species (Wojcik, 1989). Though we have compiled 38 records, there are many undescribed species within genera known to parasitize ants (Masner & Garcia R, 2002), and thus many new records can be expected as the diversity of this family is uncovered. We report eight genera of diapriid wasps parasitizing eight ant genera.

Based on the general biology of the diapriids, it appears that adults likely enter the nest independently and have evolved a suite of morphological and behavioral adaptations to facilitate acceptance (Lachaud & Pérez-Lachaud, 2012; Loiácono *et al*., 2013). It has been suggested that diapriids originally parasitized Diptera, and over the course of their evolutionary history slowly changed hosts to members of the Formicidae, particularly in the case of the army ant subfamily Ecitoninae, where many dipterans live amongst ant refuse piles ((Huggert & Masner, 1983), within (Loiácono *et al*., 2013)). Thus, we might expect a spectrum of associations, with some diapriids as nest associates of ants and others having evolved to full endoparasitoidism of ant brood. For many of the Diapriidae associated with ants, the true host-parasite relationship is unknown.

For those diapriids that are true parasitoids of ants, it seems likely that the adult wasps do not live inside the ant nests (Fernández-Marín *et al*., 2006). Fernandez-Marín et al. (2006) note that following emergence from their cocoons, diapriid wasps are very aggressively bitten by the ants. Based on colony prevalence rates found in the field, it seems that diapriid parasitism might have a more profound impact on colony demography than that of other parasitoid families (e.g. Eucharitidae, Phoridae) (Fernández-Marín *et al*., 2006; Lachaud & Pérez-Lachaud, 2012), though this hypothesis has not yet been formally examined either empirically or theoretically.

### 4.11 Diptera

The Diptera, or true flies, are a megadiverse insect order estimated to contain 1,000,000 species (Mayhew, 2007). Of these, an estimated 16,000 species of dipterans in 21 families are parasitoids (Eggleton & Belshaw, 1992; Feener Jr & Brown, 1997). Parasitoidism as a life cycle evolved over 100 times in the Diptera, from ancestors that were mostly saprophagous (Eggleton & Belshaw, 1992; Feener Jr & Brown, 1997). Interested readers should refer to both Feener and Brown (1997) and Eggleton and Belshaw (1992) where the ecology, biogeography, and evolution of diptera as parasitoids have been extensively reviewed.

We report 492 records of Diptera from six families, the Chloropidae, Ephydridae, Phoridae, Tachinidae, Helosciomyzidae and Syrphidae as parasites and parasitoids of ants (Table S2). The Phoridae are parasitoids of adult ants and very occasionally ant brood. The Tachinidae and Helosciomyzidae are typically parasites and parasitoids of other insects, but few records have been found of them infecting ants. The Syrphidae have long been known as predators living inside ant nests, and only recently have they been recorded as true primary parasites of ants.

#### 4.11.1 Chloropidae [1 record]

One record of a fly from the family Chloropidae is known to infect ants. *Pseudogaurax paratolmos* is reported infecting *Apterostigma dentigerum*. Gonzalez et al. (2016) report that *Pseudogaurax* are solitary ectoparasites of ant larvae. They note that parasitized brood were treated the same as non-infected nest mates (González *et al*., 2016).

#### 4.11.2 Ephydridae [2 records]

Two records of parasitic flies from the family Ephydridae are reported infecting ants. *Rhynchopsilopa* are ectoparasites infecting *Crematogaster sp*. The flies mount and feed on the ants and are not known to cause host mortality (Freidberg & Mathis, 1985).

#### 4.11.3 Phoridae [485 records]

The Phoridae are a large family of flies comprising many different life history strategies. Parasitoids within the Phoridae parasitize adult ant workers, and rarely, female reproductives or brood (Williams & Whitcomb, 1974; Disney, 1994; Feener Jr & Brown, 1997; Pérez-Lachaud *et al*., 2017). Many additional phorid species have been found in association with ant colonies, as scavengers of nest detritus, as predators of ant brood and injured workers (Brown, Kung, & Porras, 2015), or as other nest associates (Rettenmeyer & Akre, 1968). While a large number of phorid parasitoids of ants have been described, host records for many species remain unknown or unpublished (Brown, Schneider, & LaPolla, 2011), and associations between some phorid flies and ant hosts remain unclear, so we can expect many more records to be added in coming years. We report 485 parasite records within the Phoridae, comprising 32 genera and 262 species parasitizing 40 unique ant genera and 182 unique ant species. Most of these are primary parasitoids of adult ants, but phorids from four genera (*Aenigmatias, Apodicrania, Nothomicrodon*, and *Pseudogaurax*) are ecto- and endoparasitoids of ant brood.

For the phorids that are parasitoids of adult ants, the general life cycle is as follows. Adult female phorids use visual and olfactory cues to locate and oviposit onto adult ant workers, using recruitment trail pheromones or nest sites of hosts as possible orientation cues (Donisthorpe, 1927; Brown & Feener Jr, 1991; Feener Jr & Brown, 1997). Phorids might capitalize by ovipositing into ants injured during army ant raids (Feener Jr & Brown, 1997). Oviposition can occur in the head, thorax, or gaster of the ant hosts, after which point larval development begins (Disney, 1994). Larvae feed on the host hemolymph, and in many species, larvae migrate to the host's head, ultimately causing decapitation. The location of this decapitation and the emergence of the adult fly can happen inside the nest, but then exiting the nest before being detected and killed by ant hosts presents a challenge. It appears that some phorid flies might be able to control the behavior of their host, inducing them to leave the colony several hours prior to emergence (Henne & Johnson, 2007). The degree to which ant behavior might be manipulated by phorid parasitoids and the mechanisms underlying such manipulation are unknown.

Many phorids associated with ants are not parasitoids and it seems that parasitoidism evolved following a scavenger-host association (Feener Jr & Brown, 1997). Thus, we find a diverse array of associations between phorids and ants, many of which have not been fully elucidated.

Phorids have been investigated extensively for their potential use as biological control agents against pestiferous ant species (reviewed in (Morrison, 2000, 2012; Folgarait, 2013). The impacts of phorid parasitoids on colony functioning appears to be low, but indirect effects of phorid parasitoidism might serve to mediate interspecific competition (Brown, 1992; Fowler & di Romagnano, 1992; Porter *et al*., 1995; Morrison, 2012). In reviewing the phorid parasitoids of leaf-cutter ants, Folgarait (2013) note that the overall parasitism rate is higher than that of *Pseudacteon* parasitizing *Solenopsis*, and that prevalence likely depends on the health status of the colony (Folgarait, 2013).

#### 4.11.4 Tachinidae [2 records]

The Tachinidae are a large family of true flies that are parasites or parasitoids of other arthropods. Few records of association between tachinids and ants exist and only two of these are parasitic in nature. *Strongylogaster globula* is an endoparasite of *Lasius* queens (Gösswald, 1950). Rettenmeyer et al. (2011) report several tachinid species within the genus *Calodexia* associated with *Eciton burchelli* during swarm raids, but the nature of the relationship remains unknown (Rettenmeyer *et al*., 2011).

#### 4.11.5 Helosciomyzidae [1 record]

We report one record of flies from the family Helosciomyzidae parasitizing ants. *Helosciomyza subalpine* is a brood ectoparasite of *Monomorium antarcticum* (reported as *Chelaner antarcticus*, (Barnes, 1980)) in New Zealand. Fly larvae chew holes in the integument of the ant larvae, and feed on the liquid exudate until the ant larvae dies several days later.

#### 4.11.6 Syrphidae [1 record]

Syrphidae are a family of flies in which the adults commonly feed on plant material (nectar, pollen), while the larvae are insectivorous or saprotrophic. The known biology of the Syrphidae associated with ants is well-reviewed in Reemer (2013) and Perez-Lachaud et al. (2014) (Reemer, 2013; Pérez-Lachaud *et al*., 2014). While the syrphid subfamily Microdontinae have long been associated as myrmecophiles living in ant nests as predators of ant brood (reviewed in (Reemer, 2013)), the first truly parasitic syrphid fly was only discovered in 2014.

Perez-Lachaud et al. (2014) report the first record of a syrphid as a primary parasite of ants, *Hypselosyrphus trigonus* infecting brood of *Neoponera villosa* (reported as *Pachycondyla villosa)* in Mexico (Pérez-Lachaud *et al*., 2014). Unlike the predatory Microdontinae which lay eggs in the vicinity of the ant nest (Feener Jr & Brown, 1997), it appears adult *H. trigonus* females oviposit eggs directly on the ant brood. As noted by Perez-Lachaud et al. (2014), *N. villosa* have not been observed moving their nests, thus it seems likely that the syrphid flies must enter the ant nests independently to access and oviposit onto brood (Pérez-Lachaud *et al*., 2014).

Syrphids within the family Microdontinae are specialized predators of ant brood. Adult Microdontinae lay their eggs outside of ant nests, and these larvae enter and develop inside ant nests. The ability to live undetected inside ant colonies is no simple feat; syrphids in the genus *Microdon* are able to mimic the cuticular hydrocarbons of their ant hosts to avoid detection (Howard, Akre, & Garnett, 1990a; Howard, Stanley-Samuelson, & Akre, 1990b). Possessing the ability to mimic one's host likely necessitates host specialization for these Microdontinae (Reemer, 2013). The impact of both predatory and parasitic syrphids on host ant colony functioning is unknown, but potentially large (reviewed in (Reemer, 2013)).

### 4.12 Strepsiptera [23 records]

The Strepsiptera (‘twisted wing parasites') are a small order of insects that spend much of their lives as entomophagous parasitoids of other insects (Kathirithamby, 1989, 2008). Females in all strepsipteran families other than the Mengenellidae spend their entire lives inside their hosts, while adult males are free-living but extremely short-lived. Only strepsipterans in the family Myrmecolacidae parasitize ants.

Female strepsipterans reproduce viviparously; the first larval instars consume their mother from the inside and then emerge through her head, which protrudes from the host’s body. Following emergence, these free-living larvae then actively seek out hosts. In the case of the Myrmecolacidae, males and females exhibit sexual dimorphism in their host choice, with males parasitizing ants while females parasitize orthopterans. Having found an ant host, the male strepsipteran larvae penetrate the cuticle and once inside, go through several larval stages before pupating. This development itself does not kill the host, rather, the host must remain alive until after the strepsipterans have completed their development. When the males are ready to emerge, they leave the host via the puparium, leaving a hole in the host that often becomes colonized by fungi (Kathirithamby, 2008). Ants parasitized by strepsipterans have been observed behaving unusually (Ogloblin, 1939; Hughes, Moya-Raygoza, & Kathirithamby, 2003; Kathirithamby *et al*., 2010).

The very short lifespan of the free-living, adult male strepsipterans and the entirely endoparasitic lifestyles of adult female strepsipterans make recording host associations with these insects difficult. Though ants were first recorded being infected by strepsipterans by Westwood in 1861, knowledge of host records remains limited, perhaps due in part to the ant collecting methods often employed by myrmecologists (Kathirithamby & Hughes, 2002).

Known host associations between strepsipterans and ants are summarized (Kathirithamby & Hughes, 2002; Hughes *et al*., 2003; Cook, 2009; Kathirithamby *et al*., 2010; Kathirithamby, 2017). Many additional strepsipterans in genera known to infect ants have been described but have unknown host associations (see Strepsiptera database). We report 23 records of Strepsiptera from the genera *Caenocholax, Myrmecolax*, and *Stichotrema* infecting nine ant genera (Table S2).

#### 4.12.1 Myrmecolacidae [23 records]

The Myrmecolacidae are the only family within the Strepsiptera to use ants as hosts during their life cycle. Currently, three genera within the Myrmecolacidae are known: *Caenocholax, Myrmecolax*, and *Stichotrema*.

*Caenocholax fenyesi* is a species complex with cryptic diversity (Hayward et al 2011); our records database will grow as this diversity is unraveled and further host associations are uncovered. *Caenocholax fenyesi sensu lato* has been recorded parasitizing *Camponotus*, *Crematogaster*, *Dolichoderus*, *Myrmelachista, Pheidole*, and *Solenopsis* in North and South America.

*Myrmecolax* has been recorded parasitizing *Camponotus, Eciton*, and *Pachycondyla* in the Neotropics and Indo-Australia region.

*Stichotrema* has been recorded from *Camponotus, Crematogaster, Pheidole, Pseudomyrmex*, and *Solenopsis* in South America, Africa, and Southeast Asia.

### 4.13 Potential records bias

Bias is undoubtedly present in the records of parasitic organisms infecting ants that we have collected here (see discussion). To begin to assess where bias might be present in the reported parasite records, we plot the number of parasite records reported for a given ant genus as a function of the total number of recognized, extant species in that ant genus (Fig. 13). A simple linear regression was calculated to predict the number of parasite records for a given ant genus based on the number of extant species in that genus. The predicted number of parasite records for a given genus is equal to 0.529 + 0.952*(n. species in genus). Though a significant regression equation was found (F(1,310) = 178.6, p < 2.2 x10^−16^), the correlation between the number of parasite records and the number of extant species in a given ant genus was weak (R^2^ = 0.37). This suggests that while the number of species within a genus is probably important, other factors might also contribute to the patterns in parasite records that we observe.

**Figure 13.**
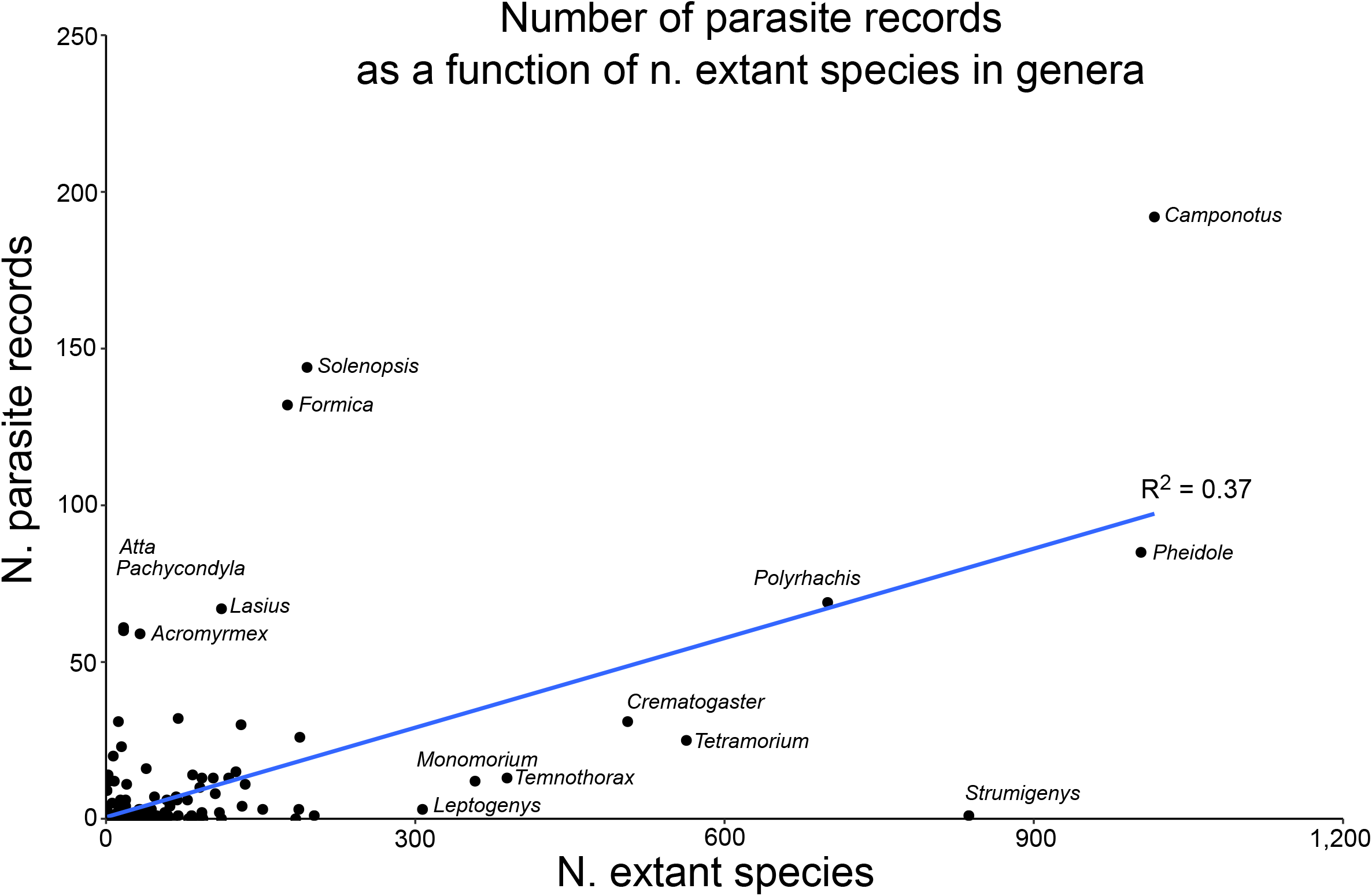
Number of parasite records as a function of species in host genus. A scatterplot showing the number of parasite records for a given ant genus on the y-axis, plotted against the number of extant species reported for that genus on the x-axis. A fit line for a simple linear regression of n. parasite records as a function of n. extant species in genus is plotted in blue, given by the formula: predicted number of parasite records for a given genus = 0.529 + 0.952*(n. species in genus).

### 4.14 Estimating the biodiversity of parasites infecting ants

A major goal of this work is to understand whether ants may be lacunae for parasite biodiversity, harboring many genera and species that have hitherto been undescribed. While it is difficult to estimate how many unobserved parasites infecting ants exist, we can use information from the collected records to make some simplistic estimates of potential ant parasite biodiversity.

We will make the simplifying assumption that ant genera without parasite records have not been adequately sampled, leaving 252 genera for which we will assume we have no parasite information. These genera have on average 14 species, and using the formula from our linear regression above, we thus predict 0.529 + 0.952(14 species per genus) = 13.857 records per genus, resulting in > 3,500 additional records that could be expected if these genera were sampled for parasites. This estimate is likely to be extremely conservative, as few targeted surveys of ant parasites have been carried out and thus the number of records for even well-studied ant genera is likely low. Given that the majority of parasite genera only infect a single host genus (Fig. 11a), we should expect many additional parasite genera and species to be discovered when we explicitly search for ant parasites.

## 5. Discussion

The goal of this work was to collect records of parasites infecting ant hosts in order to synthesize major patterns in the natural history and epidemiological relationships of these parasites with their hosts. To that end, we report 1,415 records of parasites and parasitoids in 51 families infecting 82 genera of ants (Table S2). The majority of the parasitic organisms we report infecting ants are parasitoids (89.6%, 1,268/1,415 records, Fig. 6), which require death of the ant host as a developmental necessity. Most parasitic organisms infecting ants encounter the hosts that they will subsequently infect in the extranidal (outside of the nest) environment (primary encounter-68.6%, 970/1,415 records, Fig. 7), though others are able to enter colonies independently of host behavior (independent encounter - 10.2%, 144/1,415 records, Fig. 7). To complete their life cycles, most parasitic organisms infecting ants only need to use a single host (direct transmission-89.2%, 1,262/1,415 records, Fig. 8), though the use of indirect life cycles involving multiple hosts predominates amongst the worms (Trematoda, Cestoda, and Nematoda, Fig. 8). Finally, most parasitic organisms infecting ants need to leave the nest to complete their life cycles before they are capable of transmitting to new ant hosts *(ex-nido* transmission-88.3%, 1249/1,415 records, Fig. 9). Most parasitic organisms infecting ants are endoparasites (84.1%, 1,190/1,415 records, Fig. 10). Notably, most parasites infecting ants are specialists, with each parasite genus typically infecting only one ant genus (Fig. 11).

A second major goal was to understand whether the ants, one of the most ecologically successful groups of organisms and which have a diverse array of life history strategies, are host to a diverse group of parasites. Our synthesis of parasite records confirms that ants are host to a diverse group of parasitic organisms, ranging from viruses and bacteria to helminths and insects. Of the major taxonomic groups infecting ants, Order Diptera (flies), Phylum Fungi, and Order Hymenoptera (parasitic wasps) predominate, accounting for 34.8%, 25.6%, and 25.0% of the records, respectively (Fig. 5). Viruses and bacteria account for only 1.1% of the total parasite records (16/1,415 records, Fig. 5). Some parasite groups are likely undersampled, in most cases due to lack of targeted surveys and particularly for those parasites that are difficult to identify (e.g. Strepsiptera).

However, we lack records of parasitism for the majority of ant species; less than 4% of all valid, extant ant species and subspecies have a single parasite record associated with them (580/15,334, (Bolton, 2018)). We found that less than 25% of all currently recognized ant genera have any parasite records associated with them (82/334 genera, Table S1), and fewer than 55% of extant ant sub-families have any records (9/17 sub-families, Fig. 4). Of the genera that do, the median number of reported records is five (Fig. S2), with the genera *Camponotus, Solenopsis, Formica, Pheidole*, and *Polyrhachis* dominating the records (192, 144, 132, 85, and 69 records, respectively). The number of reported records for an ant genus has a general positive correlation with the number of species in that genus, but the fit of a simple linear regression to the data is poor (R^2^ = 0.37), suggesting that many ant genera may be undersampled. Other ant genera, particularly those with invasive or pestiferous species, may have received more scrutiny in order to identify potential bio-control agents, leading to a preponderance of records for those genera.

### 5.1 Why does parasitoidism predominate?

The predominance of parasitoidism (89.5%) amongst the reported records may seem counterintuitive at first glance - killing off hosts as developmental necessity leaves fewer future hosts to infect. However, ants are eusocial organisms and their fitness lies at the level of the colony, not at the level of the individual (Wilson, 1971; Sherman *et al*., 1988). Thus, killing off one individual, or even many individuals, may have negligible impacts on lifetime colony fitness. Indeed, the flexible division of labor that is the hallmark of ant colonies (Macevicz & Oster, 1976; Hölldobler & Wilson, 1990, 2009) likely allows colonies to buffer the loss of individuals, whether it is due to disease, predation, or inter- and intraspecific competition (Hughes *et al*., 2008). Furthermore, the presence of age-related polyethism in many ant species (Hölldobler & Wilson, 1990) means that many of the individuals infected by parasites (particularly those which use primary encounter strategies) are amongst the oldest in the colony and are thus expected losses. In this light, parasitoids may simply impose an extra mortality term on these oldest age classes. Provided that this extra mortality term doesn't over-run what the colony can replace, parasitoids can benefit while colonies buffer the loss.

### 5.2 Why do most parasitic organisms of ants use *ex-nido* transmission?

*Ex-nido* transmission, in which parasitic organisms must leave the ant colony to complete development, find other hosts, or mate, is extremely common across the majority of taxa infecting ants. *Ex-nido* transmission predominates amongst the worms and insects, and it is also common amongst specialist fungal parasitoids (e.g. *Ophiocordyceps*). For parasites that use multi-host life cycles (e.g. worms), *ex-nido* transmission is a natural consequence of needing to find hosts that do not live inside ant colonies. For others, their development might require specialized microclimates that cannot be found inside nests (e.g. ‘zombie-ant' fungi, *Ophiocordyceps*, (Andersen *et al*., 2009)). But for many parasites that use direct life cycles only requiring ant hosts, why leave the bountiful and protected confines of the colony?

Organisms are generally quite good at recognizing self from non-self, and the ant colony superorganism is no exception. Ants live in a chemical world ((Hölldobler & Wilson, 1990) p. 197 - 297), and are constantly assessing nest mates for their signature colony smell via hydrocarbons that are applied across the ant's cuticle. Ants can quickly discriminate nest mates from non-nest mates, and experiments have shown that nest mates that have been covered in the hydrocarbons of other colonies are shown aggression (Bonavita-Cougourdan, Clément, & Lange, 1987; Morel & Vander Meer, 1987). Accordingly, ant colonies are typically on high alert for any potential intruders. For parasitic organisms that utilize free-living stages in their life cycle (e.g. adult flies, worms), rapid detection once they are outside of their ant host is likely. Thus, it becomes imperative for these organisms to get outside of the colony quickly. For parasitic organisms that do use *in-nido* transmission, the strong recognition systems of colonies can act as a further barrier to within-nest disease transmission. Infected individuals can be recognized and removed from colonies (Ugelvig *et al*., 2010; Pull *et al*., 2018). The adoption of *ex-nido* transmission strategies is therefore either a biological necessity on the part of the parasite, or likely a consequence of strong anti-intruder defenses by ant colonies.

For some parasites that require extended periods of development inside their ant host, avoiding detection as ‘non-self' is critical. These parasites need their infected ant hosts to remain inside the nest and continue receiving food and to that end, may have evolved the ability to mask their own smell whilst inside their hosts. For example, it has recently been shown that ants infected by the ‘zombie ant' fungi *Ophiocordyceps*, which can take weeks to grow before it kills its host, are not discriminated against by their non-infected nest mates (Solá Gracia *et al*., 2018), even though they are filled with fungi as they near death (Hughes *et al*., 2011; Fredericksen *et al*., 2017). Social parasites and associates that live extended lives inside ant nests also need to avoid host detection (reviewed in (Lenoir *et al*., 2001)). Other parasitic organisms may exploit the sophistication of the colony recognition system for their own aims by manipulating the cuticular hydrocarbons of their infected hosts over time. For example, nematodes infecting ants require a long period of development inside their ant host; however, they also need to access other host species in aquatic environments. To this end, they may abruptly change the cuticular hydrocarbon profile of their infected host so that nest mates treat the ant aggressively, forcing it out of the colony (D. Hughes, unpublished data), and getting it one step closer to where its next transmission event will occur.

### 5.3 Patterns in the records of parasitic organisms infecting ant genera

The observed patterns in parasitic organisms infecting ants are the result of three major factors, discussed below: (i) ant and parasite ecologies, (ii) ant and parasite evolutionary and co-evolutionary histories, and (iii) biases in how records have been collected.

#### 5.3.1 Ant and parasite ecologies

The association of ants and parasites depends on both their evolutionary past (below) as well as their ecological present. Certain ant nesting or foraging ecologies may pre-dispose them to associations with certain parasites or parasitoids. For example, nesting on or in soil or leaf litter may pre-dispose ant species to infections by generalist fungi, whereas we may expect to find fewer records of generalist fungi infecting arboreal species (Boomsma, Schmid-Hempel, & Hughes, 2005). We may expect to find more infections of brood in ant species that are nomadic or polydomous, as in these situations brood are moved around and might be more exposed to parasitic organisms that infect using primary encounters. For ant species that tend hemipterans for their honeydew, we might expect to find instances where viruses or bacteria known to infect hemipterans have made host jumps into ants. Indeed some ant colonies nesting near bees have been found harboring viruses that typically infect bees (Gruber *et al*., 2017). Machine learning techniques (e.g. boosted regression trees, (Elith *et al*., 2006) have been used to correlate host life history traits with parasite records for other animal groups (Han *et al*., 2015). Future work could apply this methodology to understand which ant traits correlate to observed parasite records, and to predict parasite-host combinations for which we have not yet found association records.

#### 5.3.2 Ant and parasite evolutionary histories

The evolutionary pasts of both ants and their parasites have played a role in determining their extant relationships reported here. Though a comprehensive treatment of how parasite records assort across the ant phylogeny is outside the scope of this current work, it could be helpful to lay the future groundwork for this using some general concepts borrowed from the theory of island biogeography (MacArthur & Wilson, 1967). Ant sub-families and genera can be imagined as host islands colonized by parasites. As the ants diversified, these host islands may have shrunk (e.g. become more specialized in their ecological niche, changed habitats or geographic range), presenting a challenge to their parasites. Not all of the parasites may have made the transition alongside their host and for those that did, host ecological specialization may have fostered subsequent parasite specialization or diversification. Additionally, throughout the host diversification process, new parasite families, genera, and species were arising or coming into contact with these ant host islands, providing opportunities for new colonization events.

While it will be difficult to tease apart the full evolutionary history of ants and their parasites, the rich phylogeny of the ants (Moreau *et al*., 2006; Moreau & Bell, 2013) coupled with that of some of their parasites (e.g. eucharitid wasps, Murrary and Heraty 2013), will make uncovering parts of this story possible. For example, Murray and Heraty (2013) found that eucharitid wasps, which exclusively parasitize ants, made hosts shifts early in their evolutionary history, but then maintained host conservatism even after dispersal events and speciation (Murray *et al*., 2013). Additionally, they found congruence in the phylogenies of ants and eucharitids, suggesting that their evolutionary histories are more similar than random chance alone would predict. Hopefully, as more associations between ants and their parasites become known and as phylogenies become better resolved, we can start piecing together the larger evolutionary story of ants and their parasites.

#### 5.3.3 Biases in parasite records

There is no doubt that bias exists in the parasite records that we report here. Bias exists in which parasitic organisms have been the most extensively studied, and indeed, our knowledge of many particular parasite groups is due to one or several individuals who have championed those taxa and made studying them their life's work. There remain many additional parasite taxa infecting ants that are in dire need of natural historians to take up their cause. Our understanding of the relationships between ants and their parasites has also been hindered by observational bias; it is much easier to notice larger parasites, such as worms and insects, than microscopic parasites (e.g. viruses, bacteria) that have only recently become culturable or identifiable due to methodological advances. One case in point is that of viruses- the first virus known to infect ants was first identified in 2004, though ants are likely host to many viruses (Valles *et al*., 2004). As we begin surveying for parasites and parasitoids more explicitly, more records of ant-parasite association will undoubtedly be discovered.

Bias is also present in the ant genera that have been most studied. Certain ant genera have ecologies that make them more amenable to study. For example, even though the genus *Strumigenys* is quite speciose (837 species, (Bolton, 2018)), their small size and cryptic nesting ecologies make them more difficult to study generally, and thus we know far less about their general biology and their interactions with parasites. In contrast, other ant genera are infamous for their roles as invasive or pestiferous species (e.g. fire ants, leaf cutter ants, crazy ants, argentine ants) and they have been actively surveyed for natural enemies that could help control their populations. Accordingly, for these genera, we have far more reported parasite records.

### 5.4 Open questions, future directions, and a call to action

In this work, we have assembled known records and synthesized major patterns in the life history traits of parasites infecting ants. This provides a first step towards approaching the many open questions and important areas of research that remain, some of which we detail here. Firstly, we still lack knowledge of how individual loss due to disease impacts colony level survival and fitness, or how this impacts colonies at different stages of development (i.e. incipient or mature colonies). We don't know how intensely colonies experience parasite pressure, whether certain parasites pre-dominate in exerting that pressure, and how that parasite pressure scales with latitude or other biogeographic factors. We have limited understanding of inter-colony and metapopulation parasite transmission dynamics. We also have little understanding of how parasites of ants impact the larger communities in which they reside. In answering these questions, we will be better positioned to understand how sociality impacts parasite diversity, how parasite diversity impacts the evolution of sociality, and how parasite virulence evolves with social hosts.

We hope that this works serves as a call to action, to myrmecologists, to parasitologists, and to evolutionary ecologists. In order to understand the ecological success of the ants, we need to understand the relationships that they have had with their parasites, both now and over evolutionary time. To further appreciate parasite diversity and understand the roles of parasites in ecological communities, we need to understand the relationships that they have with their hosts. Finally, the ants, with their well-resolved phylogeny and diversity of host ecological traits, offer an exciting opportunity for testing theories of evolutionary ecology, and present a comparative foil to other social living groups.

## 6. Conclusions

1. Ants are predominantly infected by parasitic flies (Order Diptera), fungi (Phylum Fungi), and wasps (Order Hymenoptera), with very few records of viruses and bacteria known to cause morbidity or mortality.
2. The majority of parasitic organisms infecting ants are parasitoids, requiring host death as developmental necessity, and most directly encounter their hosts outside of the nest.
3. Most parasitic organisms infecting ants require a period of time outside of the nest before they are capable of transmission to future ant hosts, preventing direct transmission between ants inside the nest.
4. Less than 4% of extant ant species and fewer than 25% of currently recognized ant genera have any recorded parasites. We conservatively estimate that we are missing at least 3,500 additional records, many of which will represent previously undescribed parasite genera and species.

## 7. Acknowledgments

We gratefully acknowledge the many natural historians who have collected records of parasites infecting ants and other social insects. Their hard, often underappreciated, work provides the foundation upon which all of our biological knowledge is built. We would like to thank Jessica M. Conway, Andrew F. Read, and Peter J. Hudson for comments and helpful feedback on earlier versions of this work. This material is based upon work supported by the National Science Foundation Graduate Research Fellowship under Grant No. DGE1255832 to L.E.Q., NIH Grant R01 GM116927-02 to D.P.H., and NSF Grant No. 1414296 as part of the joint NSF-NIH-USDA Ecology and Evolution of Infectious Diseases program. Any opinion, findings, and conclusions or recommendations expressed in this material are those of the authors(s) and do not necessarily reflect the views of the National Science Foundation.

